# Pre-epidemic evolution of the USA300 clade and a molecular key for classification

**DOI:** 10.1101/2022.12.05.519169

**Authors:** Colleen Bianco, Ahmed M. Moustafa, Kelsey O’Brien, Michael Martin, Timothy D. Read, Barry Kreiswirth, Paul J. Planet

## Abstract

USA300 has remained the dominant community and healthcare associated methicillin-resistant *Staphylococcus aureus* (MRSA) clone in the United States and in northern South America for at least the past 20 years. In this time, it has experienced epidemic spread in both of these locations. However, its pre-epidemic evolutionary history and origins are incompletely understood. Large sequencing databases, such as NCBI, PATRIC, and Staphopia, contain clues to the early evolution of USA300 in the form of sequenced genomes of USA300 isolates that are representative of lineages that diverged prior to the establishment of the South American (SAE) and North American (NAE) epidemics. In addition, historical isolates collected prior to the emergence of epidemics can help reconstruct early events in the history of this lineage. Here, we take advantage of the accrued, publicly available data, as well as two newly sequenced pre-epidemic historical isolates from 1996, and a very early diverging ACME-negative NAE genome to understand the pre-epidemic evolution of USA300. We use database mining techniques to emphasize genomes similar to pre-epidemic isolates, with the goal of reconstructing the early molecular evolution of the USA300 lineage. Phylogenetic analysis with these genomes confirms that the North American Epidemic and South American Epidemic USA300 lineages diverged from a most recent common ancestor around 1970 with high confidence, and it also pinpoints the independent acquisition events of the of the ACME and COMER loci with greater precision than in previous studies. We solidify evidence for a North American origin of the USA300 lineage and identify multiple introductions of USA300 into South America from North America. Notably, we describe a third major USA300 clade (the pre-epidemic branching clade; PEB1) consisting of both MSSA and MRSA isolates circulating around the world that diverged from the USA300 lineage prior to the establishment of the South American and North American epidemics. We present a detailed analysis of specific sequence characteristics of each of the major clades, and present diagnostic positions that can be used to classify new genomes.

## Introduction

*Staphylococcus aureus* is a major cause of human disease worldwide. Clonal complex 8 (CC8) is one of the most successful *S. aureus* lineages and has given rise to several major methicillin-resistant *S. aureus* (MRSA) clones, the most prominent of which is the USA300 clone that emerged as the dominant cause of community-associated (CA) infections in the USA (Seybold et al., 2006, King et al., 2006). USA300 is a virulent clone that was first identified in the United States during an outbreak of infections starting in November 1999 in a Mississippi state prison (2001). Prior to the 1990s, most MRSA infections were associated with hospital settings, but the USA300 clone rapidly became widespread in the general population around the year 2000, and by 2004, it had become the major cause of severe soft-tissue infections in the United States and the dominant CA-MRSA circulating in North America (King et al., 2006). The prototypical USA300 clone is characterized by key genetic features: point mutations in genes *cap5D* and *cap5E* resulting in a lack of a functional capsular polysaccharide (Mohamed et al., 2019, Boyle-Vavra et al., 2015), the pathogenicity island SaPI5 encoding the enterotoxin genes *sek* and *seq*, Panton-Valentine leukocidin (PVL) encoded by genes *lukF-PV* and *lukS-PV*, possession of the staphylococcal chromosomal cassette mec IVa (SCC*mec*IVa), and, most uniquely, the locus referred to as the arginine catabolic mobile element (ACME) (Diep et al., 2006, Kennedy et al., 2008), which appears to have been acquired by horizontal gene transfer just prior to the spread of the epidemic (Planet et al., 2013).

After USA300 was identified in the United States, a closely related USA300 substrain was increasingly detected in northern South American countries (Alvarez et al., 2006). The first reported isolate of this CA-MRSA lineage was isolated in 2005 in Colombia and this lineage was soon noted to be spreading through community and hospital settings in Colombia, Venezuela, and Ecuador (Alvarez et al., 2006, Arias et al., 2008). This USA300 Latin American variant (USA300-LV) appeared to cause the same spectrum of disease as USA300 from North America, and it had many of the key genetic signatures of the North American USA300 lineage, including a similar pulsed field gel electrophoretic (PFGE) pattern, possession of SaPI5, *lukF-PV*, and *lukS-PV* (Arias et al., 2008). USA300 isolates from South America were found to differ from North American USA300 in two key molecular features: they mostly contained a SCC*mec*IVc locus and a mobile genetic element with genes conferring copper and mercury resistance (COMER) in place of ACME (Arias et al., 2008, Planet et al., 2015, Reyes et al., 2009). Although there is a close relationship between isolates considered to be USA300-LV and those from North America, all USA300-LV isolates appeared to have diverged prior to the beginning of the North American epidemic with a major clade of isolates from the South American Epidemic (SAE) forming the immediate sister clade to the North American Epidemic clade (NAE) (Planet et al., 2015). Molecular clock estimates suggest that NAE and SAE lineages shared a common ancestor 40-50 years ago with each epidemic emerging independently in parallel in the 1980s for SAE and early 1990s for NAE (Planet et al., 2015, Strauß et al., 2017, Copin et al., 2019, Uhlemann et al., 2014).

Although analyses have favored a North American origin for all USA300 lineages, the geographic origin of the common ancestor of SAE and NAE has not been solidly established (Planet et al., 2015, Strauß et al., 2017). The early branching (EB) lineages of the USA300 clade that diverged prior to the establishment of the North American and South American epidemics, are made up of isolates from both geographical regions and are very sparsely sampled (Planet et al., 2015). This observation suggests that *S. aureus* USA300 was likely circulating at low levels in both regions prior to the establishment of the epidemic clades, and it may have experienced multiple introductions from one region to the other (Planet et al., 2015). EB lineages lack the clear genetic hallmarks from each epidemic clade (COMER from SAE and ACME from NAE (Planet et al., 2015)) and the genomic features that unite them with the epidemic lineages, and likewise distinguish them, are incompletely characterized.

Understanding the origins and evolution of these early branching isolates is critical to understanding the emergence of both epidemics.

A huge increase in the numbers of available *S. aureus* genomes presents an opportunity to revisit the pre-epidemic evolution of USA300 and identify factors that led to the success of the epidemic lineages. However, identifying which genomes will be informative, from the tens of thousands available, can be difficult. We used the comparative genomic tools WhatsGNU (Moustafa and Planet, 2020) and PATRIC (Davis et al., 2019) to identify and obtain genomes with a specific focus on the EB genomes from the USA300 tree. We also sequenced and added 2 historical isolates collected in 1996, which, through molecular-typing appeared to be close relatives of the USA300 clade (Roberts et al., 1998). Our analysis confirms that the NAE and SAE USA300 clades diverged from a most recent common ancestor around 1970 (95%HPD 1966-1974). We solidify evidence supporting a North American origin for both the NAE and SAE clades, and we identify a large clade made up of both MSSA and MRSA isolates with a worldwide distribution that diverged prior to the establishment of the South and North American epidemics, referred to here as PEB1. We also present diagnostic sequence changes in the early evolution of the USA300 clade that can be used both as a classification tool (a molecular key) and to understand possible biological changes that led to the success of USA300 and its sublineages.

## Methods

### Whole Genome Sequencing

Whole genome sequencing was performed for isolates 2m-n, 65-669, BK2651 and BK2448. Genomic DNA was prepared using either the DNeasy Blood and Tissue kit (Qiagen) or the Wizard Genomic DNA Purification Kit (Promega) after lysostaphin treatment. Genomic DNA libraries for 2m-n, 65-669, and BK2448 were prepped using the NexteraXT DNA sample preparation kit and sequenced on a HiSeq sequencer (Illumina) with 250-bp paired-end reads. Genome assembly was done using Unicycler pipeline (Wick et al., 2017). The complete genome of BK2651 was determined using Oxford Nanopore Technology (MinION) and Illumina MiSeq sequencing. MinION sequencing libraries were prepared using the rapid barcoding kit and sequenced using a MinION flow cell. The Unicycler pipeline was used for hybrid de novo assembly of Illumina and MinION reads. Sample metadata and genomes were deposited in GenBank under the following BioSample identifiers: 2m-n: SAMN31430433; BK2448: SAMN31431371; BK2561: SAMN31431372; 65-669: SAMN10689409. Genome quality statistics are shown in Table S1.

### Finding similar genomes

We used the similar genome finder utility of WhatsGNU (Moustafa and Planet, 2020) to find the 100 closest genomes in the Staphopia database (Version: 06/27/2019, contains 43,914 genomes, (Petit and Read, 2018)) to the following isolates chosen to represent basal portions of the USA300 tree: 65-669 (GenBank assembly accession: GCA_016107225.1), BK2651 (Roberts et al., 1998), M121 (Planet et al., 2015), V2200 (Planet et al., 2015). We also used the Similar Genome Finder Service tool to find 50 similar public genomes in PATRIC (Davis et al., 2019) based on genome distance estimation using Mash/MinHash. PATRIC utilizes a database drawn largely from NCBI consisting of 27,000 genomes. The resulting lists of 150 genomes most similar to each query genome were combined and duplicates were removed, resulting in 204 genomes. These genomes were screened for sequencing contamination using MASH (Ondov et al., 2019), resulting in 198 genomes. All genomes are freely available on NCBI and individual accession numbers are listed in Supplemental excel file S1.

### Phylogenetic analysis

A maximum likelihood tree was constructed for 276 genomes; 198 genomes from the closest genome screen, 70 genomes associated with the North and South American epidemics (Planet et al., 2015, Von Dach et al., 2016, Planet et al., 2016) including two reported here for the first time: 65_669 and BK2651, and 8 non-CC8 outgroups, including 2m-n and BK2448, reported here for the first time. To construct the maximum likelihood tree, reads were first trimmed using TrimGalore as follows: trim_galore --illumina -q 15 --stringency 7 --paired read1.fastq read2.fastq. Reads were mapped to the TCH1516 reference genome (GenBank accession GCF_000017095.1) and variant calling was performed using Snippy v.4.6.0 (Page et al., 2016) as follows: snippy --cpus 16 --outdir mysnps --ref tch1516full.gb --R1 Read1.fastq.gz --R2 Read2.fastq.gz, then: snippy-core --prefix core mysnps. The SNP alignment produced by Snippy was used to infer an initial phylogenetic tree in RAxML v8.2.4 (Stamatakis, 2014) as follows: raxml -T 8 -m GTRGAMMA -f d -N 1 -s core.full.aln -n Output -p 1977. The initial ML newick tree produced by RAxML and the whole-genome alignment produced by Snippy were used as input for ClonalFrameML to infer recombination (Didelot and Wilson, 2015) as follows: ClonalFrameML newick_file aln_file output_prefix. Maskrc-svg (Kwong, 2019) was used to mask the recombinant regions in the whole-genome alignment (produced by Snippy) based on the output analysis of ClonalFrameML. This new whole-genome alignment was then used to construct a final phylogenetic tree in RAxML and node support was evaluated with 100 nonparametric bootstrap pseudoreplicates using the following parameters: raxml -T 8 -m GTRGAMMA -N 100 -s alnfile -p 1977 -b 1977. The tree was visualized in iToL (Letunic and Bork, 2021). The maximum likelihood tree containing the 39 additional CC8 genomes as outgroups was constructed beginning with reads the same way as described above.

### Staphopia tree

We inferred a phylogenetic tree of 42,949 *Staphylococcus aureus* whole genomes sequences by first calculating pairwise MASH distances between them. RapidNJ (Martin Simonsen, 2008) (https://birc.au.dk/software/rapidnj) was used to infer a phylogenetic tree from these pairwise distances. The tree was rooted at the longest branch prior to visualization in ggtree (Yu, 2020) and ggplot2 (Villanueva and Chen, 2019). Sequence type annotations for each tip in the tree were taken from the Staphopia database (Petit and Read, 2018).

### Divergence time estimation

To estimate the emergence time of the clades in the tree, we used a branch tip calibrated approach using a Bayesian phylogenetic framework implemented in BEAST v2.6.0 (Bouckaert et al., 2019). We used the whole-genome alignment produced using Maskrc-svg that accounts for recombination (see above). The SNP-sites tool was used to extract SNPs from this alignment (Page et al., 2016). The SNP alignment was then used to estimate divergence times in BEAST (Bouckaert et al., 2019). The Hasegawa–Kishino–Yano (HKY) nucleotide substitution model was used with estimated base frequencies (Hasegawa et al., 1985). Because a SNP alignment was used instead of a whole genome alignment, ascertainment bias for variable-only sites was corrected for by editing the XML file to factor in the number of invariant sites based on fully sequenced genomes (https://www.beast2.org/2019/07/18/ascertainment-correction.html). We ran three analyses; one strict clock as described below and two relaxed clocks as described in Figs. S1-S2. We implemented a strict clock model with a random starting tree and a coalescent constant population using 300 million Markov chain Monte Carlo (MCMC) steps with a 5,000-step thinning. After the 10% of the first posterior samples were removed as a burn-in, the MCMC trace determined the effective sample size values to be above 110 for all parameters and the maximum clade credibility tree was determined using TreeAnnotator v2.6.3. A median rate of 1.217×10^−6^ (95% HPD, 1.1184×10^−6^, 1.3117×10^−6^) was estimated for this analysis (Fig. S3).

### Ancestral state geographical reconstruction

We used PastML (Ishikawa et al., 2019) to re-construct geographical states of common ancestors throughout the tree. We provided PastML with the rooted phylogenetic tree made in RAxML with tips annotated with geographic location. Ancestral character reconstruction was performed using MPPA+F81 model.

### Presence of mobile genetic elements

Genome reads were assembled with Unicycler (Wick et al., 2017) and annotated with Prokka (Seemann, 2014). Blast (Johnson et al., 2008), command: blastn –db nt –query nt.fasta -evalue 1e-6 –out results.out, was used to determine presence of: COMER, ACME, SapI5, and Panton Valentine Leukocidin genes according to nucleotide sequences in USA300_FPR3757 (GenBank assembly accession: GCA_000013465.1). The whole COMER sequence from CA15 (Planet et al., 2015) and the whole ACME sequence from FPR3757 were used as query sequences. Query sequences of SapI5 (*sek* and *seq*) and PVL (*lukF* and *lukS*) were obtained from FPR3757. SCC*mec* finder (Kaya et al., 2018) was used to determine SCC*mec* type.

### Screen for genetic markers of pre-epidemic USA300 evolution

Roary (Page et al., 2015) and Scoary (Brynildsrud et al., 2016) were used to identify genes uniquely present or absent in each clade. We used annotated assemblies in GFF3 format produced by Prokka (Seemann, 2014) to calculate the pangenome in Roary. The output from Roary was used by Scoary to determine statically significant genes found in each clade compared to the other clades. WhatsGNU (Moustafa and Planet, 2020) was used to identify gene protein alleles specific to each clade and a python script (unpublished, A. Moustafa) was used to sort alleles according to clade and execute statistical tests (sensitivity, specificity, p-value, odds-ratio, Bonferroni correction) similar to Roary. WhatsGNU was used to identify genomes in Staphopia with exact matches to the diagnostic alleles (-i, --ids_hits option).

To identify diagnostic SNPs we used Mesquite (Maddison and Maddison, 2021) ancestral character reconstruction to identify synapomorphies that uniquely characterized each major clade. Character reconstruction was performed on a maximum parsimony (MP) consensus tree generated in PAUP 4.0a (Feb 10 2021 Build) (Swofford, 2003) using the SNP matrix. The MP tree was determined using a tree-branch reconnection (TBR) heuristic search with all characters unordered and given equal weight. All character state transitions were weighted equally. The consensus tree was nearly identical to the RAxML tree. SNPs predicted to be changing on the branch leading to each clade were determined first. Then each of these positions was further assessed for the following criteria: (1) appearance of either the ancestral or derived nucleotide in all isolates in the tree, (2) 100% concordance (presence) of the derived SNP in every member of the clade, (3) 100% concordance (presence) of the ancestral SNP in the outgroup.

## Results & Discussion

To better understand the early evolution and origins of the USA300 epidemics, we sampled genomic databases for genomes similar to four *S. aureus* genomes that represented USA300 early-diverging lineages. We used the similar genome finder utility of WhatsGNU to query the Staphopia database (43,914 genomes). WhatsGNU identifies genomes that have the highest numbers of exact protein allele matches to the query genome. We also used the Similar Genome Finder Service tool to find similar genomes in PATRIC (Davis et al., 2019) which utilizes a database drawn largely from NCBI and consists of 27,000 genomes. To query these databases, we used two well-characterized genomes that fell clearly outside of the NAE clade: the SAE isolate M121 and the EB isolate V2200. We also used two new genomes that we are reporting here for the first time: an early diverging NAE isolate 65-669 (isolated in New York in 2012 as part of a collection of *S. aureus* isolates from atopic dermatitis patients; Project PRJNA512846) and the EB isolate BK2651 (isolated in New York in 1996 (Roberts et al., 1998)). Preliminary phylogenomic analysis had suggested that BK2651 was the earliest isolate known that was clearly in the USA300 clade. The isolate 65-669 was chosen because, in preliminary analysis, it appeared to be the earliest diverging member of the NAE clade known to date.

In total, out of 600 hits (150 for each query genome), 204 unique genomes were identified from the Staphopia and PATRIC databases. The large number of NAE genomes represented in this analysis along with a high percentage of overlap amongst “best hit” sets, suggests that we were able to find most of the EB and SAE genomes in the database. The 204 genomes were isolated between 1999 and 2017 and originated in Africa (n = 3), Australia (n = 2), Europe (n = 21), North America (n = 129), Asia (n = 1), and South America (n=23). Twenty-five genomes had no geographical information. For the phylogenetic analyses, we included 64 well-characterized NAE, SAE or EB isolates (Planet et al., 2015, Von Dach et al., 2016, Planet et al., 2016) and 8 outgroup genomes, including two additional genomes: 2m-n and BK2448, reported here for the first time (Roberts et al., 1998) (Table S1).

SNP-matrices were used to reconstruct maximum-likelihood (Fig. 1) and Bayesian (Fig. 2) trees. Based on the tree topologies, our analysis included 130 NAE, 73 SAE, and 65 genomes that branched before the emergence of these 2 clades (not including known outgroup genomes). Following previous notation, these 65 genomes would be considered “Early Branching” genomes (Planet et al., 2015) because they are the early branches of the USA300 tree. Thirty-six of these genomes made up a prominent, well-supported (Fig. 1; bootstrap value of 100) early branching lineage that we designate here as the pre-epidemic branching USA300 clade 1 (PEB1).

**Figure 1:**
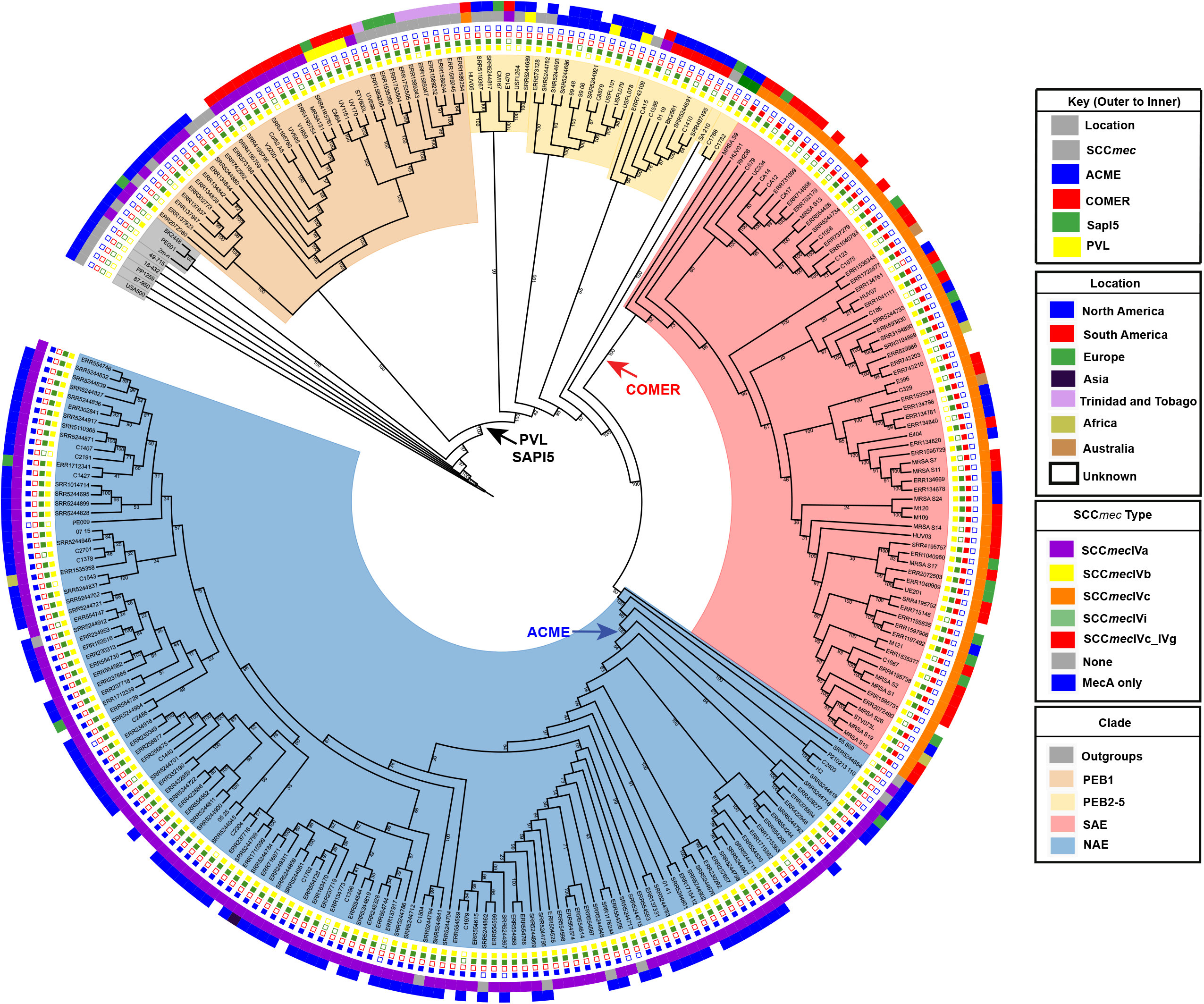
Maximum likelihood tree of 276 *S. aureus* genomes: Isolation location (outer track) and SCC*mec* type (inner track) is shown for each genome. Presence or absence of key genetic elements are indicated with solid or hollow squares. Introduction of key genetic elements are indicated. White squares indicate missing data.

**Figure 2:**
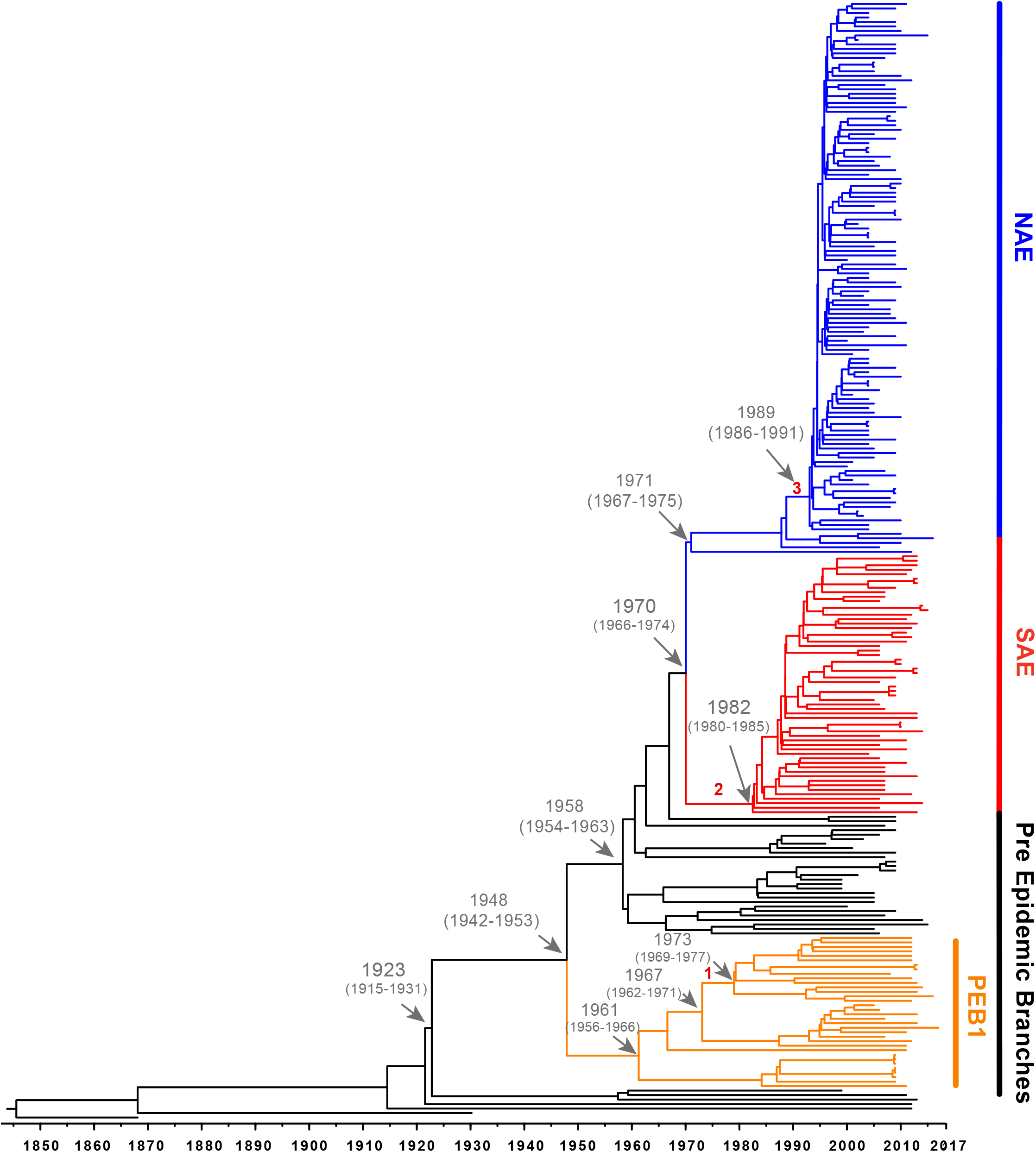
Bayesian maximum clade tree calculated from 54001 sampled trees. Tree generated using a strict clock and a constant-size coalescent population. Important MRCA are indicated as dates (95% HPD) and introduction of key genetic elements are labelled (1. vSaß loss: 1967-1973 (95% HPD 1962-1977), 2. COMER acquisition: 1970-1982 (95% HPD 1966-1985), 3. ACME acquisition: 1988-1989 (95%HPD 1985-1991.)

### The Early Branches of the USA300 clade

Likely reflective of higher rates of sampling in North America, genomes belonging to the NAE clade were overrepresented in our dataset accounting for almost one-half (130/276, 47%) of genomes. Almost all of these genomes were isolated in North America. In line with previous reports, the majority of these isolates were PVL+, SapI5+, ACME+ and carried SCC*mec*IVa. We also identified five ACME-negative NAE genomes SRR5244854, P210213_110, C2403, H2, and 65-669. Previous analyses have noted ACME negative NAE isolates (Strauß et al., 2017, Uhlemann et al., 2014, Copin et al., 2019) that are usually thought to represent loss of the mobile element. However, some NAE isolates lacking ACME (eg., C2403 (Planet et al., 2015) isolated in the USA in 2010) have been shown to branch at the base of the NAE clade likely representing lineages of this clade that diverged before ACME was acquired (Planet et al., 2015, Strauß et al., 2017). For the ACME-negative genomes identified here, SRR5244854 was isolated in 2006 in Illinois (USA), P210213_110 was isolated from the sputum of a Tennessee (USA) man in 2010, H2 was isolated in 2016 from an Austrian river (Lepuschitz et al., 2018), and 65-669 was isolated in 2012 in a study of atopic dermatitis. Based on their position on the tree these 5 isolates are representative of the ancestor of NAE before ACME was acquired. Focusing on the branch separating these early diverging NAE lineages from the rest of the clade, molecular clock analysis suggests that ACME was acquired between 1988-1989 (95%HPD 1985-1991), which is consistent with previous estimates timing the acquisition of ACME in the late 1980s (Planet et al., 2015, Strauß et al., 2017). Of the five NAE ACME-negative genomes in our tree, three possess SCC*mec*IVa while two genomes, 65-669 and P210213_110 are MSSA, indicating, along with the presence of SCC*mec*IVa in other early branching lineages, that ACME and SCC*mec*IVa were likely not acquired together.

Using the SAE genome M121 as bait, we were able to identify 41 new SAE isolates. In agreement with previous reports, these genomes were PVL+, ACME-negative, COMER+ and carried SCC*mec*IVc (Planet et al., 2015). Our results support previous work that SAE acquired SCC*mec*IVc and the COMER element between 1970-1982 (95% HPD 1966-1985) (Planet et al., 2015). While most of the isolates in SAE are from South America (n=33; Columbia, Ecuador, and Venezuela) there are also genomes from Europe (n=11, Demark, Germany), Australia (n=2), and North American (n=13, United States), indicating transmission of SAE to these locations. Our analyses support a most recent common ancestor for the two epidemic lineages in 1970 [95% HPD 1966-1974].

The EB lineages of the USA300 tree are of particular interest because the origins of these early branching isolates could be key to understanding the origin of the SAE and NAE common ancestor. Due to sparser sampling, the topology of this portion of the tree has been less robust. With the added genomes from our database search, we were able to identify 5 pre-epidemic lineages with robust bootstrap support (Fig. 1). The largest of these, and the earliest to diverge, is the PEB1 clade noted above. This clade features 36 isolates that form three subclades corresponding to geographic isolation location: North America, South America and Trinidad and Tobago/Germany. According to our molecular clock analysis, this clade emerged in 1948 (95% HPD 1942-1953) (Fig. 2) after the acquisition of PVL and SAPI5 by the most recent common ancestor of the entire USA300 clade. All isolates from PEB1 are ACME and COMER-negative, and they contain SCC*mec* IVa or IVb. No mobile elements were observed in the genomic location occupied by ACME or COMER. PEB1 contains three genomes, V2200, MRSA131, and V1859, that were originally included in the description of USA300-LV (Arias et al., 2008).

The other 4 pre-epidemic branching clades together contained 29 genomes. Isolates from these clades are mostly PVL-positive (n=25), MRSA (n = 21) and MSSA (n = 8) from North America (n = 20), South America (n = 3), and Europe (n = 2). Interestingly, the MRSA genomes that make up the pre-epidemic branching clades contain various SCC*mec* types (IVa n=19; IVb n=7; IVc n=1; IVi n=2), supporting the notion of multiple introductions of SCC*mec* into USA300 (Strauß et al., 2017).

The tree also allowed us to update our understanding of the geographic origins of pre-epidemic USA300 (Fig. 3). We used PastML (Ishikawa et al., 2019) and Mesquite (Maddison and Maddison, 2021) to re-construct geographical states of common ancestors. The ancestral reconstruction definitively reconstructs the main trunk of the tree that gives rise to the PEB lineages and the two epidemic clades as being in North America, further solidifying a likely North American origin for this lineage. Because this reconstructed origin, there are five putative introduction events of USA300 from North America in the pre-epidemic part of the tree into South America (Fig. 3). Interestingly, pre-epidemic branching genomes isolated from South America have SCC*mec*IVa and those from North America have SCC*mec*IVc, in direct contrast to the SAE and NAE which have SCC*mec*IVc and SCC*mec*IVa, respectively (Planet et al., 2015).

**Figure 3:**
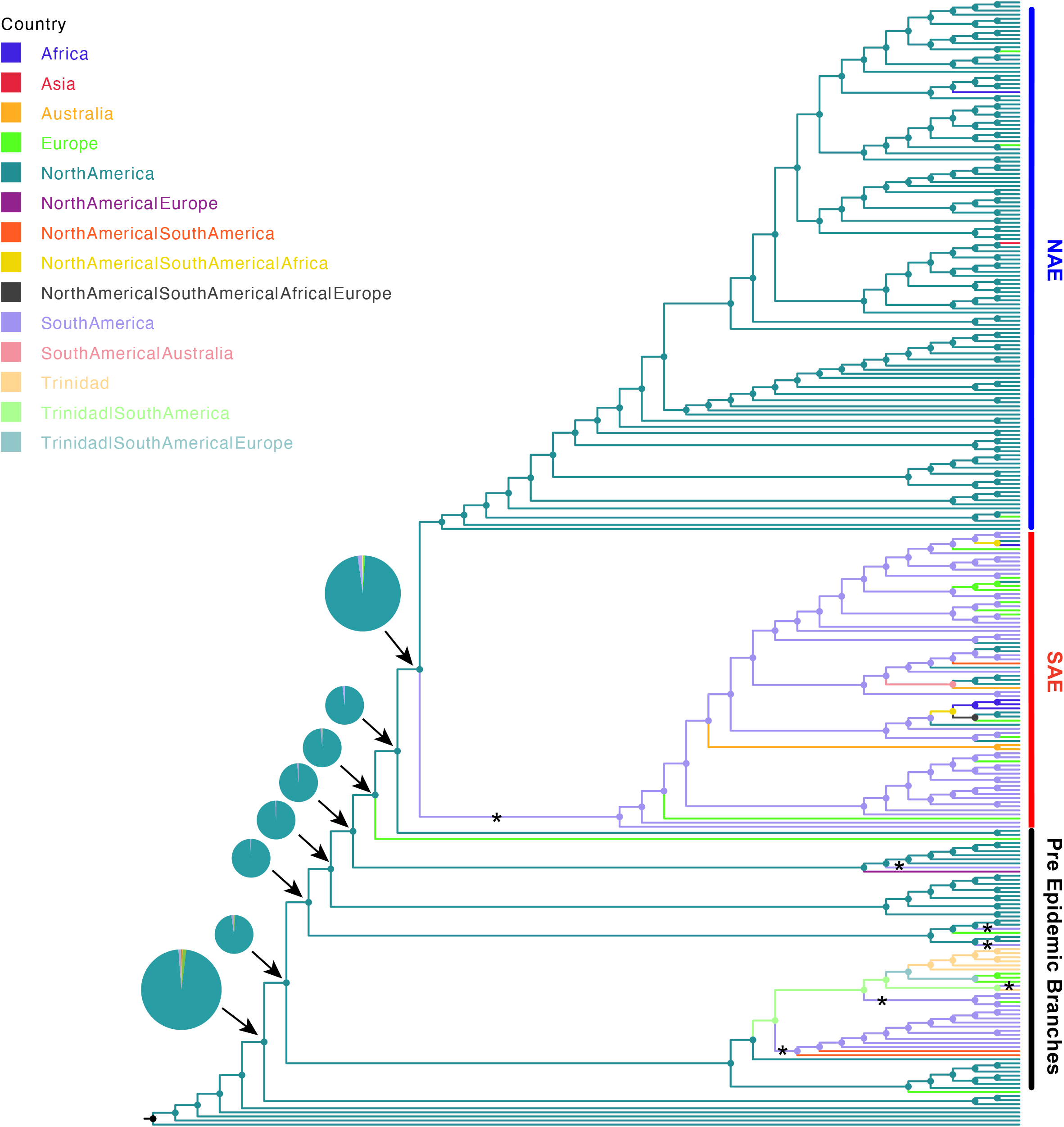
Ancestral state reconstruction of *S. aureus* isolates. Ancestral state reconstruction was done using log likelihood restricted maximum a posteriori analysis of the phylogenetic tree with the classification of each sequence based on collection location. Branches are colored according to the most probable location state of their descendant nodes as indicated at the legend. “*” indicates introduction from North America to South America. The pie charts show the probability of location for each common ancestor on the major branch leading to the NAE and SAE clades.

### Evolution of capsular polysaccharide genes

Unlike many other *S. aureus* lineages, USA300 does not produce a surface capsular polysaccharide (CP) due to mutations in the *capABCDEF* operon (Boyle-Vavra et al., 2015). Common mutations (compared to the CP+ *S. aureus* Newman) include a frameshift mutation in a polyadenine (AA) tract in the *cap5D* gene resulting in a truncated Cap5D enzyme (*cap5D*_994AA) and a point mutation (*cap5E*-223) in *cap5E* that converts Asp to Tyr in the enzyme active site resulting in an inactive protein (Boyle-Vavra et al., 2015) (Table S2). Previous analysis of these *cap5* mutations in USA300 and USA500 *S. aureus* genome sequences suggested that the *cap5D* insertion (*cap5D*_994AA) occurred in the last common ancestor of all USA300 and USA500 genomes whereas the *cap5E*-223 mutation had its origins in an ancestor of all USA300 and was acquired after the split with USA500 (Boyle-Vavra et al., 2015). Additionally, it has been reported that NAE acquired the *cap5E*-223 mutation, simultaneously with ACME and SCC*mec*IVa, in the late 1980s (Strauß et al., 2017).

To understand the acquisition of the *cap5* mutations, we evaluated the appearance of the *cap5* mutations on our *S. aureus* tree (Fig. 4). In agreement with previous studies, the majority of USA300 isolates in our tree contained the *cap5D* insertion (*cap5D*_994AA) mutation. Surprisingly, eight isolates contain a WT *cap5D*, five of which form a subclade in PEB1, one is in another PEB lineage, and two of which are found in the SAE clade. All of these isolates also contain the WT *cap5E* allele and no other mutations in the *capABCDEF* operon, indicating that these isolates likely do form a capsule. This is consistent with previous reports noting clades within USA500 where this mutation reverted to WT (Boyle-Vavra et al., 2015). This is the first report of WT *cap5D* in USA300. Of note, the WT *cap5D* allele was found in BK2561 which was isolated in 1996, the earliest USA300 genome known to date. Seven of the eight genomes with the WT *cap5D* allele were isolated in the United States between 1996 and 2009, which may explain anecdotal reports of USA300 expressing capsule.

**Figure 4:**
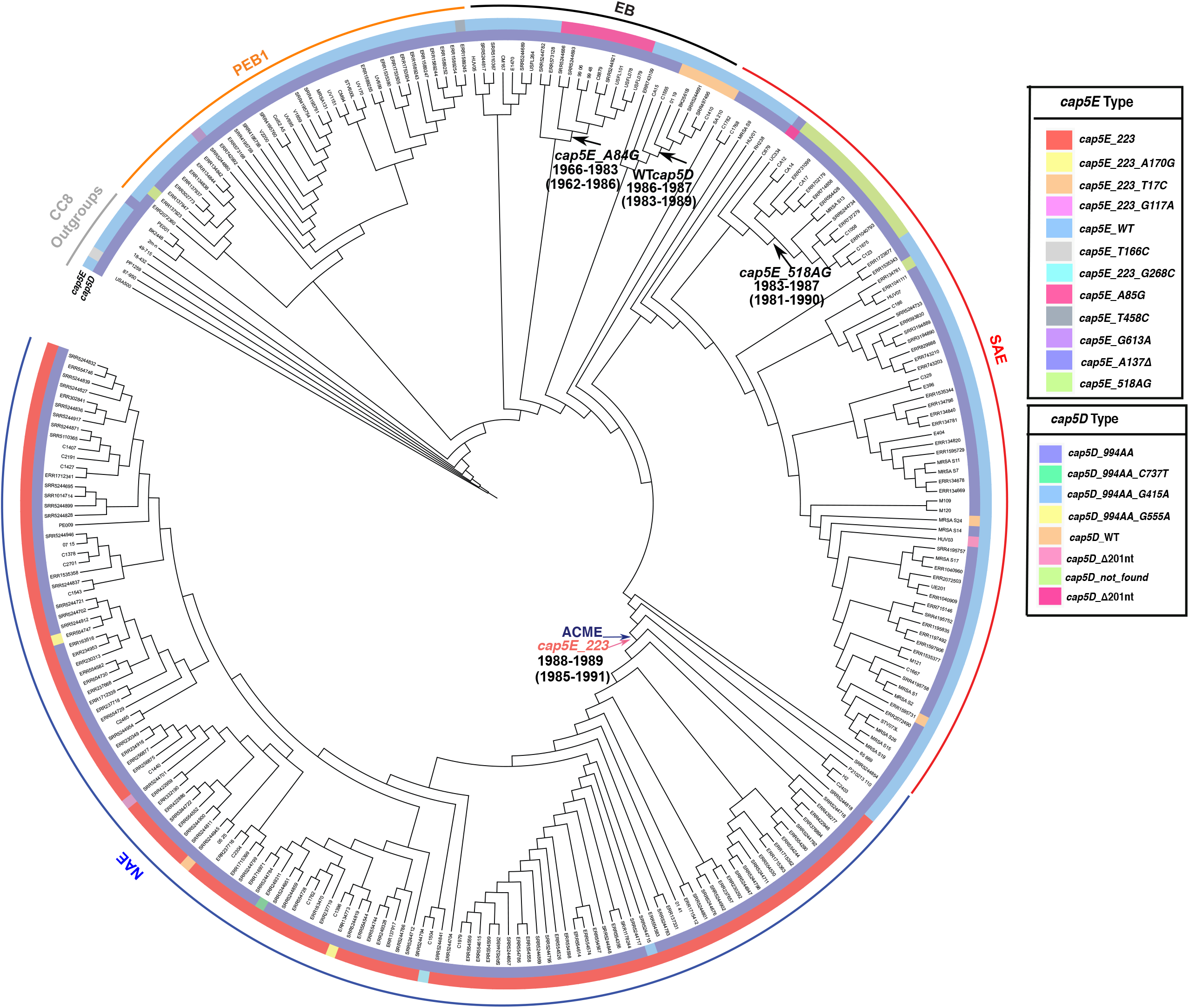
Phylogenetic analysis of *S. aureus* strains showing the distribution of conserved mutations in *cap5* in USA300: The *cap5D* (inner track) and *cap5E* (outer track) allele type is shown for each genome. Introduction of ACME and *cap* mutations are indicated.

Our analysis further shows that the *cap5E*-223 allele is present only in the NAE clade while SAE and the pre-epidemic branching genomes mostly possess WT *cap5E*. While the majority of NAE isolates contain this mutation, the ACME-negative, basal NAE isolates 65-669, SRR5244854, P210213_110, H2, and C2403 contain WT *cap5E*. As noted previously, these five isolates diverged prior to the addition of ACME within the NAE clade. This is concordant with a near simultaneous acquisition of this mutation with the introduction of ACME (Fig. 4).

### Identification of Diagnostic Genes, Protein Alleles, and SNPs

Given our well supported tree, we sought to identify sequence characteristics that were diagnostic for each of the major clades. Such genomic features can serve two important purposes. First, they offer a low-cost classification system without the need for extensive phylogenetic analysis. Second, they may give insight into the initial functional changes that happened in the evolution of the USA300 clade. We looked for diagnostic genomic features at the following three levels: (1) gene presence/absence, (2) protein alleles, and (3) SNPs. To assess gene presence absence we used Roary (Page et al., 2015) and Scoary (Brynildsrud et al., 2016). To determine diagnostic protein alleles we used the exact protein match tool WhatsGNU (Moustafa and Planet, 2020), and to identify diagnostic SNPs we used an ancestral reconstruction technique as implemented in Mesquite (Maddison and Maddison, 2021). Our goal was to identify features that were as specific as possible such that the feature was never seen outside the group, and then secondarily as sensitive as possible so that we required a minimum number of markers to classify a genome.

### Identification of markers for pre-epidemic branching isolates and PEB1

Roary/Scoary (supplemental excel file 2) identified no whole gene presence or absence that was 100% sensitive and specific for the PEB1 clade, meaning that there were no genes that offered a clear marker. Notably, we found that 27 of the 36 genomes in the PEB1 clade were missing *splABCDEF*, an operon of serine proteases found in *Staphylococcus aureus* Genomic Island *v*Saβ (Fig. S4). The genomic island *v*Saβ harbors a number of virulence-associated genes, such as the lantibiotic gene cluster (bacteriocin *bsa*), two leukocidin genes (*lukD* and *lukE*), and a cluster of serine protease genes (serine protease like, *spl*, genes) as well as two genes belonging to the type I staphylococcal restriction-modification (RM) system (*hsdM* and *hsdS*). Upon closer observation, we noted that in addition to missing the *splABCDEF* operon, these genomes are also missing the majority of *v*Saβ. Typically, *v*Saβ encodes about 30 genes. The PEB1 genomes that are missing *splABCDEF* only encode eight of these genes; seven genes found at the 3’ end of *v*Saβ (*epiABC, lukD, lukE*, and two genes that encode hypothetical proteins) and one lipoprotein, encoded at the very 5’ end of *v*Saβ (Fig. S4). The *v*Saβ integration site is well conserved among strains of *S. aureus*, with a tRNA cluster at the 3’ end and genes for two hypothetical proteins at the 5’ end, both of which are consistently present in the genomes that lack most of *v*Saβ (Fig. S4A).

The *v*Saβ genomic island integration was likely mediated by a phage, followed by diversification (through multiple recombination, integration, and excision events) into the types observed today (Kläui et al., 2019). Based on this, we hypothesize that *ν*Saβ was present in the ancestral *S. aureus* USA300 lineage, acquired through a phage integration event, with subsequent loss of part of the island in one sublineage of PEB1 isolates. Our molecular clock analysis dates this excision event between 1967-1973 (95% HPD 1962-1977) (Fig. S4B). It should be noted that four other genomes in other clades in this tree are also missing this part of *v*Saβ, indicating that this excision event is not uncommon. These genomes are sporadically distributed in the tree and did not form a distinct branch as observed in the PEB1 clade.

In contrast to the gene presence/absence approach, WhatsGNU did identify a protein allele in PEB1 (Table 1 and supplemental excel file 3) that was 100% specific and 100% sensitive for the PEB1 clade. This protein allele was a variant of HisB (imidazoleglycerol-phosphate dehydratase). To understand the physiological implications of the amino acid changes (Ala53Gly) in this version of HisB (Table 1), we used an in-silico approach. We used the program SIFT: Sorting Intolerant From Tolerant (Ng and Henikoff, 2003), which predicts whether an amino acid substitution affects protein function based on sequence similarity and the physical properties of amino acids (Table S3). The Ala53Gly change was predicted to not be functional. We also used PoPMuSiC (Dehouck et al., 2011), which evaluates the changes in folding free energy of a given protein under point mutations on the basis of the experimental protein structure. PoPMuSiC confirmed that the HisB Ala53Gly mutation would be expected to destabilize the thermodynamic and thermal stability of the enzyme.

**Table 1:** Unique SNPs and protein alleles of USA-300 strains for identification purposes. The chromosomal location of the SNP in the reference TCH1516 genome is presented in paratheses. A detailed list of nucleotide change and protein function is given in supplemental excel file 2.

|  | USA300-NAE | USA300-SAE | USA300-epidemic (NAE-SAE) | USA300-PEB1 | USA-300 |
| --- | --- | --- | --- | --- | --- |
| WhatsGNU Protein alleles | MarR, YefM, PdhD, USA300HOU_0586 USA300HOU_1426 | USA300HOU_2198, vwb | Der_2, DesR, NagE, SdcS, ComEC | HisB | LplA1, AdhR, PchA, HisG |
| SNPs | *recN* (1638449), *leuS* (1888283) | *USA300HOU_0191* (202764), *intergenic* (265666), *USA300HOU_0397* (424978), *argS* (670365), *intergenic* (835434), *USA300HOU_0795* (850349) *vwb* (876702), *USA300HOU_0938* (982790), *oppD1* (990291), *ebh* (1488257), *rluB* (1611873), *comGA* (1657533), *alaS* (1723795), *tyrS* (1843158), *USA300HOU_1746* 1877551), *intergenic* (1978879), *leuA2* (2171680), *USA300HOU_2076* (2202788), *atpA* (2224500), *USA300HOU_2105* (2233258), *USA300HOU_2169* (2310703), *USA300HOU_2198* (2341385), *intergenic* (2362258), *USA300HOU_2243* (2383283), *moeA* (2392932), *intergenic* (2405549), *USA300HOU_2322* (2459187), *USA300HOU_2384* (2525989), *USA300HOU_2423* (2561857), *USA300HOU_2474* (2611197), *crtN* (2703257), *intergenic* (2727910) | *intergenic* (201647), *USA300HOU_1266* (1352385), *ebh* (1483188), *engA* (1544604), *comEC* (1700414), *USA300HOU_1715* (1839546), *USA300HOU_1918* (2063960), *intergenic* (2262458), *lacE* (2329451), *USA300HOU_2279* (2414655), *USA300HOU_2654* (2811819) | *galE* (148897), *intergenic* (191588), *pbuX* (437253), *USA300HOU_0421* (445172), *USA300HOU_0421* (445217), *set21* (452762), *thiD* (638407), *mnhD1* (697737), *intergenic* (737821), *USA300HOU_0775* (826842), *USA300HOU_0937* (982232), *abH2* (986803), *oppB3* (997946) *USA300HOU_1022* (1089485), *isdE* (1141794), *cfxE* (1235660), *dnaE1* (1284408), *miaB* (1319259), *sbcC* O1377590), *intergenic* (1414153), *gcvPB* (1650510) *hemN* (1693959), *intergenic* (1729030), *accD* (1809323), *USA300HOU_1731* (1862944), *intergenic* (1896103), *USA300HOU_1865* (2015160), *USA300HOU_2026* (2143039), *intergenic* (2203597), *USA300HOU_2279* (2414447), *USA300HOU_2283* (2418359), *USA300HOU_2482* (2619836), *intergenic* (2736965), *intergenic* (2773434), *hisB* (2836831) | *intergenic* (14363), *USA300HOU_0113* (115190), *gatC1* (291025), *USA300HOU_0276* (311262), *lplA1* (381083), *USA300HOU_0470* (495436), *USA300HOU_0498* (542064), *tagX* (714370), *USA300HOU_0721* (771214), *USA300HOU_0721* (771429), *intergenic* (810108), *intergenic* (856156), *mnhD2* (944473), *oppA1* (993238), *menF* (1047527), *pdhB* (1104230), *intergenic* (1199330), *USA300HOU_1599* (1706793), *USA300HOU_1731* (1863834), *intergenic* (2503985), *USA300HOU_2464* (2603502) |

Imidazoleglycerol-phosphate dehydratase (encoded by *hisB*) catalyzes the sixth step in the histidine biosynthesis pathway. It has been shown to play a crucial role in biofilm formation in *Staphylococcus xylosus* as well as being a potential target of the antibiotic cefquinome (Zhou et al., 2018). Deletion of *hisB* results in histidine auxotrophy in the fungal pathogens *Aspergillus fumigatus* (Dietl et al., 2016) and *E. coli* (Patrick et al., 2007), however, a *hisB* mutant in *S. aureus* has not been studied.

We used a phylogenetically informed process in Mesquite (Maddison and Maddison, 2021) to find signature SNPs at the node representing the closest PEB1 ancestor. Importantly, this analysis allowed us to also detect synonymous and intergenic SNPs. We identified 35 SNPs present in all genomes and unique to the PEB1 clade (Table 1 and Supplemental excel file 4). The SNP found in *hisB* corresponded to the nonsynonymous change identified in our WhatsGNU screen. Together with the novel alleles found using WhatsGNU, the multiple SNPs unique to the PEB1 clade give strong support to the assertion that this is a new, defined group.

### Identification of markers for epidemic USA300, NAE and SAE

We also sought to define unique genetic markers for isolates belonging to USA300 epidemic clades; SAE and NAE individually and together. Roary/Scoary analysis for presence and absence showed that genes constituting ACME were specific to NAE and that genes constituting COMER were only found in SAE as reported previously [10]. However, none of these genes were 100% sensitive for either clade with the most sensitive genes from these regions obtaining only 93% sensitivity for NAE or SAE. When grouped together the clade composed of NAE and SAE also had multiple genes with 100% specificity, but limited sensitivity. Of note, the two proteins with the highest sensitivity were CopX(B)(sens. 92%) and CopL(YdhK)(sens. 89%), which have been noted previously as the only two genes shared between ACME and COMER (Planet et al., 2015).

WhatsGNU analysis (Table 1 and supplemental excel file 3) was also able to identify alleles that are 100% specific for the NAE or SAE clades individually, but none of these alleles had 100% sensitivity. However, the highest sensitivity obtained for 100% specific protein alleles was 98% for both SAE and NAE, suggesting that an exact match protein allele approach may be a highly effective classifier with two or more genes. To build a compound classification scheme we identified combinations of unique alleles that can be used to screen for SAE or NAE clade members (Table 1). Of the numerous novel alleles specific to NAE, we identified four that were found in 128/130 (98% sens/100% spec) of NAE genomes and not found anywhere else on the tree. However, the genomes 65-669 and H2 that are basal members of the NAE clade do not contain any of these 4 alleles. The 65-669 genome along with 125 other NAE genomes do contain a novel allele for an uncharacterized lipoprotein (USA300HOU_1426) that is 100% specific to the NAE clade. Adding these alleles to the other 4 alleles, we can identify all NAE clade members except H2. We were unable to find a unique protein allele that could be used to link the H2 genomes to the NAE genome.

The SAE clade contains numerous, specific, novel protein alleles. Of these specific alleles, we identified 1 protein allele (an uncharacterized M23 family peptide) found in 72/73 (98% sens/100% specificity) SAE genomes and not found anywhere else on the tree. The ERR715146 genome did not contain this protein allele, however, this genome along with 68 other SAE genomes contained a novel staphylocoagulase allele (encoded by *vwb*) that is 100% specific and 94% sensitive for the SAE clade. Together, these two alleles can be used to identify all SAE clade genomes in our tree.

We also sought to identify diagnostic alleles within the combined NAE and SAE clade. It should be noted that these markers are also candidates for genes that may have been instrumental in the fitness of this epidemic clones. As with other groups in this tree, there were many alleles that had 100% specificity. The most sensitive 5 of the 100% specific alleles had between 92-96% sensitivity for these clades, corresponding to 7-15 genomes in the clade having a different allele (Table 1). These alleles are: GTPase Der (*der_2*), Transcriptional regulatory protein DesR (*desR*), PTS system N-acetylglucosamine-specific EIICBA component (*nagE*), Sodium-dependent dicarboxylate transporter SdcS (*sdcS*), and ComE operon protein 3 (*comEC*).

We used SIFT (Ng and Henikoff, 2003) to predict whether or not these epidemic alleles are active (Table S4). The epidemic SdcS and ComE alleles are predicted to not be active enzymes. The epidemic ComEC allele is of interest because of its involvement in natural transformation (Pimentel and Zhang, 2018). While natural transformation is a key component in the evolution of microbial populations, it remains an open question whether *S. aureus* natural competence is a frequent event or only very rare event in *S. aureus* populations (Morikawa et al., 2012). However, it was recently reported that induction of natural competence in *S. aureus* not only allows for DNA update from the environment, but also adapts staphylococcal metabolism to infection conditions by increasing the rate of glycolysis (Cordero et al., 2022), which could have impacted fitness of the epidemic strains. SdcS is a Na+/dicarboxylate symporter that transports succinate, fumarate and malate into the cell, which then feeds into the TCA cycle (Hall and Pajor, 2007). SdcS has not been extensively studied in *S. aureus*.

While it is unclear what consequences inactivation of SdcS and ComEC may have during infection, the connection to metabolic activities is intriguing. *S. aureus* undergoes substantial metabolic adaptation, especially by selective use of the tricarboxylic acid cycle, during infection (Acker et al., 2019, Bosi et al., 2016, Gabryszewski et al., 2019). Metabolism of fumarate and malate is a critical component of staphylococcal adaptation as evidenced by large increases in the expression of *fumC*, which codes for fumarate hydratase a key enzyme in the TCA cycle interconverting fumarate and malate (Acker et al., 2019, Gabryszewski et al., 2019).

The remaining epidemic specific alleles, *der_2, desR*, and *nagE*, are predicted to produce active enzymes, so it is unclear what direct effect these mutations have on the success of the epidemic lineages. Der (double Era-like GTPase) is a GTPase that plays an essential role in the late steps of ribosome biogenesis (Hwang et al., 2012). 50S subunits assembled in the absence of Der are defective and unable to assemble into 70S ribosomes, a lethal event. Der is highly ubiquitous in most bacteria and is not found in eukaryotes, making it an excellent antibiotic target candidate (Hwang et al., 2012). Based on our estimates of divergence times this mutation occurred between 1967-1970 (HPD 95% 1963-1974), just before the divergence of the two epidemic lineages, highlighting this allele as a possible target for future phenotypic study. DesR is the response regulator in a two-component system, along with the histidine kinase DesK, neither of which has been characterized in *S. aureus. S. aureus* DesK expressed in *B. subtilis* can functionally complement the *B. subtilis* homologue DesK (Fernández et al., 2020). DesKR in *B. subtilis* is involved in temperature sensing but it is unknown if this is the role of DesKR in *S. aureus* (Fernández et al., 2019). Lastly, NagE has been found to be a factor involved in human endothelial cell damage (Xiao et al., 2022). Specifically, a *nagE* mutant in the *S. aureus* JE2 (USA300 NAE) background caused significantly less damage to human epithelia compared to wild type JE2. It is tempting to speculate that this particular epidemic-specific protein sequence might cause more endothelial damage than other versions of NagE.

As above, we also used Mesquite to find SNPs acquired along the single ancestral branch of each of these clades (Table 1 and supplemental excel file 4). We found 32 SAE-specific unique genetic markers, 2 NAE-specific unique genetic markers, and 11 genetic markers diagnostic for both NAE and SAE. The SNPs unique to NAE are particularly important because we were unable to find 100% sensitive markers at the whole gene or protein allele levels. We identified two SNPs present in all genomes and unique to the NAE clade (Table 1 and Supplemental excel file 4). These SNPs were acquired between 1970-1971 (HPD 95%, 1966 to 1975), approximately 19 years before ACME was acquired. One of these SNPs, G105A (gene nucleotide location), is in the coding region of *recN*. This is a synonymous nucleotide substitution, with both codons coding for lysine. The other SNP (C12T, gene location) is located in the coding region of *leuS*, encoding the leucine-tRNA ligase, and is also a synonymous nucleotide substitution, with both codons coding for tyrosine.

### Identification of markers for the entire USA300 clade

We next sought to identify overall USA300-specific unique genetic markers. As with our previous analyses we first sought to identify whole gene differences that were diagnostic for all genomes from the USA300 taxa in our tree. Because there were only 8 non-USA300 genomes in our tree, we surmised that our techniques could incorrectly identify unique genes, alleles, and SNPs that were not specific to USA300. Thus, for this analysis we added 39 additional non-USA300 CC8 genomes derived from a previous analysis by Bowers et al (Bowers et al., 2018) as a comparison (Fig. S5). Despite many genes with 100% specificity, the highest sensitivity identified by Roary/Scoary was 96.6% for one gene; USA300HOU_0815 (encoding a hypothetical protein). Some of the Scoary-identified 100% specific genes also appeared to be homologous to similar genes found outside of USA300 (eg., LukD and HlgC) and were probably identified by procedures in Roary/Scoary for separation of orthologues and paralogues. These genes are unlikely to be useful for classification.

Our WhatsGNU analysis identified multiple alleles that are 100% specific to the entire USA300 clade, with the highest sensitivity being 98%. We were able to identify a combination of four protein alleles that classified all of the genomes in our USA300 clade. (Table 1; sequence in supplemental text file 3). One of these alleles (lipoate protein ligase 2) is present in 263/268 genomes (98% sens/ 100% spec). The three other alleles, when used in combination with the novel lipoate protein ligase 2 allele, can identify four of the remaining USA300 genomes. We were unable to find a diagnostic allele to include the fifth genome (H2), which may be due to sequence quality of this genome.

At the SNP level we also used Mesquite to find diagnostic SNPs acquired along the single ancestral branch from our non-USA300 CC8 outgroups to USA300 (Fig. S5). We found 21 USA300-specific unique genetic markers (Table 1 and supplemental excel file 4). Any or all of these SNPs can be used to determine if a newly found *S. aureus* isolate is a member of the USA300 clade.

### Application and Testing of USA300 unique clade markers

As mentioned above, diagnostic alleles were often more sensitive than whole gene presence/absence for classification. In addition, genes acquired by horizontal gene transfer may make presence/absence strategies prone to misclassification. Exact match protein alleles are much less likely to be found in other parts of the tree because of the strict criteria for defining an exact match (100% identity and 100% coverage). While protein alleles may theoretically be horizontally transferred as well, they are less likely to remain exactly the same after transfer. For these reasons we pursued a protein allele strategy to for classification. Our suggested strategy for classifying an unknown isolate as USA300 is shown in Fig. S6. Diagnostic allele sequences are listed in supplemental excel file 3.

The numbers of genomes in public databases that belong to the USA300 clade and subclades is not known, and the large numbers of genomes make phylogenetic classification computationally difficult. To test our classification strategy, we used the Staphopia database (a collection of 43,914 curated genomes). Our strategy was to use the diagnostic alleles to query this database, and then confirm the identity of these genomes using a phylogenetic approach. When querying Staphopia for USA300 genomes, we required each detected genome to have at least one of the four diagnostic alleles. Using these criteria, we identified 4097 potential USA300 genomes. We removed 27 redundant genomes and mapped the remaining 4070 genomes onto a preliminary tree of all Staphopia genomes calculated using a Mash-based Neighbor-joining approach (Fig. S7). Almost all of these genomes mapped to a clade with other known USA300 genomes. This clade also contained 163 potential false negative USA300 genomes, which includes a clade of 141 genomes and 22 other genomes scattered through this clade. Further, 9 potential false positive genomes were found outside of this clade.

We suspected that the very large NJ Staphopia tree may have some errors in it due to the computational challenges associated with very large datasets. To assess whether the potential misclassification of these genomes we added all of the misclassified genomes to our prior tree and recalculated our maximum likelihood analysis (Fig. S8). Using this tree, we confirmed that the clade of 141 genomes does not fall in the USA300 clade, and 15 of the 22 other false negatives also fell outside the USA300 clade. Six of the seven remaining false negatives were PEB with the remaining one being a NAE genome. Eight of the nine potential false positives were true positives. Based on these values, we determined the specificity and sensitivity of our USA300 diagnostic alleles to be 100% and 99.8%, respectively (Table S5A).

We also used the diagnostic alleles for PEB1, SAE, and NAE to query the Staphopia database. The single protein allele for PEB1 yielded 29 genomes that were all identified by our initial search strategy and appear in our tree (Fig. 1) as well as on the Staphopia tree in their respective clade (Fig. S7). The two protein alleles diagnostic for SAE yielded 45 genomes, 44 of which appear in our tree (Fig. 1) and 43 appear in a clade together on the Staphopia tree (Fig. S7). Two genomes (SRR4195755 and ERR134761) identified as belonging to SAE using our molecular key appeared in the NAE clade on the Staphopia tree (Fig, S7), however, were confirmed to be SAE by our USA300 tree (Fig. 1 and Fig. S8).

When querying Staphopia for NAE genomes, we required each identified genome to have at least one of the five diagnostic alleles, resulting in 3995 potential NAE genomes. All but 30 of these genomes were confirmed to belong to the NAE clade on the Staphopia tree (Fig. S6). Six of these 30 genomes did not fall in the USA300 clade on the Staphopia tree, however, were confirmed to be USA300-NAE in our additional phylogenetic analysis (Fig. S8). There were 26 USA300-NAE genomes in the Staphopia tree classified as USA300 but not NAE by our molecular key. We determined that all of these genomes were pre-epidemic branching lineages (Fig. S7). The specificity and sensitivity of our NAE diagnostic alleles for this data was 99.9% and 99.4%, respectively (Table S5B).

As an additional test we applied our classification scheme to a USA300 genome, 2003-0063 (also known as SRR5244961), that has previously been of uncertain status and was not identified for our initial analysis presented here (Bowers et al., 2018). This genome was previously placed in the combined SAE and NAE clade (Bowers et al., 2018). This genome has all the novel protein alleles defined by WhatsGNU that characterize USA300 overall (lipoate protein ligase 2, AdhR, PchA, and HisG). However, the 2003-0063 genome did not contain any novel alleles defined for SAE, NAE, or PEB1. It did contain the 5 alleles specific to the combined NAE and SAE clades, supporting previous analysis placing it as a part of the SAE/NAE clade but not part of either SAE or NAE. However, this genome had the two diagnostic SNPs for the NAE group. To confirm these results, we performed further phylogenetic analysis including 2003-0063 (Fig. S5), which places it as a very early branching NAE member that diverged prior to the acquisition of ACME. Combined with the results above, this finding suggests that the diagnostic SNPs for NAE will be important additional tools for identifying very early diverging members of NAE.

### Report of historical isolate genomes sequenced

In this paper we report two historical USA300 isolates: BK2651 and BK2448. These isolates were collected in May of 1996 in New York City as part of a hospital surveillance program (Roberts et al., 1998). Isolates in this study were previously characterized using molecular typing techniques, mainly pulsed-field gel electrophoresis. BK2651 and BK2448 were both reported to be SCC*mec*IV, PVL- and ACME-. Based on whole genome sequencing, we determined that BK2448 contains SCC*mec*IVa and is PVL-, SapI5-, ACME-, and COMER-. On our tree (Fig. 1), BK2448 is part of an outgroup to USA300 with the genomes 2m-n and PE001, which branched prior to the acquisition of PVL and the most recent common ancestor the USA300 clade. BK2651, on the other hand, diverged prior to the North and South American epidemics, but within the USA300 clade. To our knowledge BK2651 the oldest, USA300 sequenced to date. Another close relative of USA300 has been reported, though it was not sequenced, that was isolated in 1995, a year prior to the isolation of BK2651, but this isolate was ACME- and PVL-, suggesting that it may have not fallen within the USA300 clade (David et al., 2015). BK2651 is PVL+, SapI5+. The SCC*mec* type for BK2651 was inconclusive, being either SCC*mec*IVg, based on similarity to individual genes, or SCC*mec*IVc, based on similarity to the whole SCC*mec* cassette.

#### Evolutionary Scenario

Combined, the observations presented here suggest the following evolutionary scenario (Fig. 5): The most recent common ancestor of USA300 was present in North America in the 1940s and had already acquired genes for the Panton Valentine Leukocidin and the SaPI5 locus at some point in the past 30-40 years. This ancestor also already had the *cap5D* mutation making it unable to make capsule. This ancestor was probably also a methicillin resistant strain with SCC*mec*IVa as the most likely cassette type, but the heterogeneity of SCC*mec* types in the early evolution of the clade makes this conclusion uncertain.

**Figure 5:**
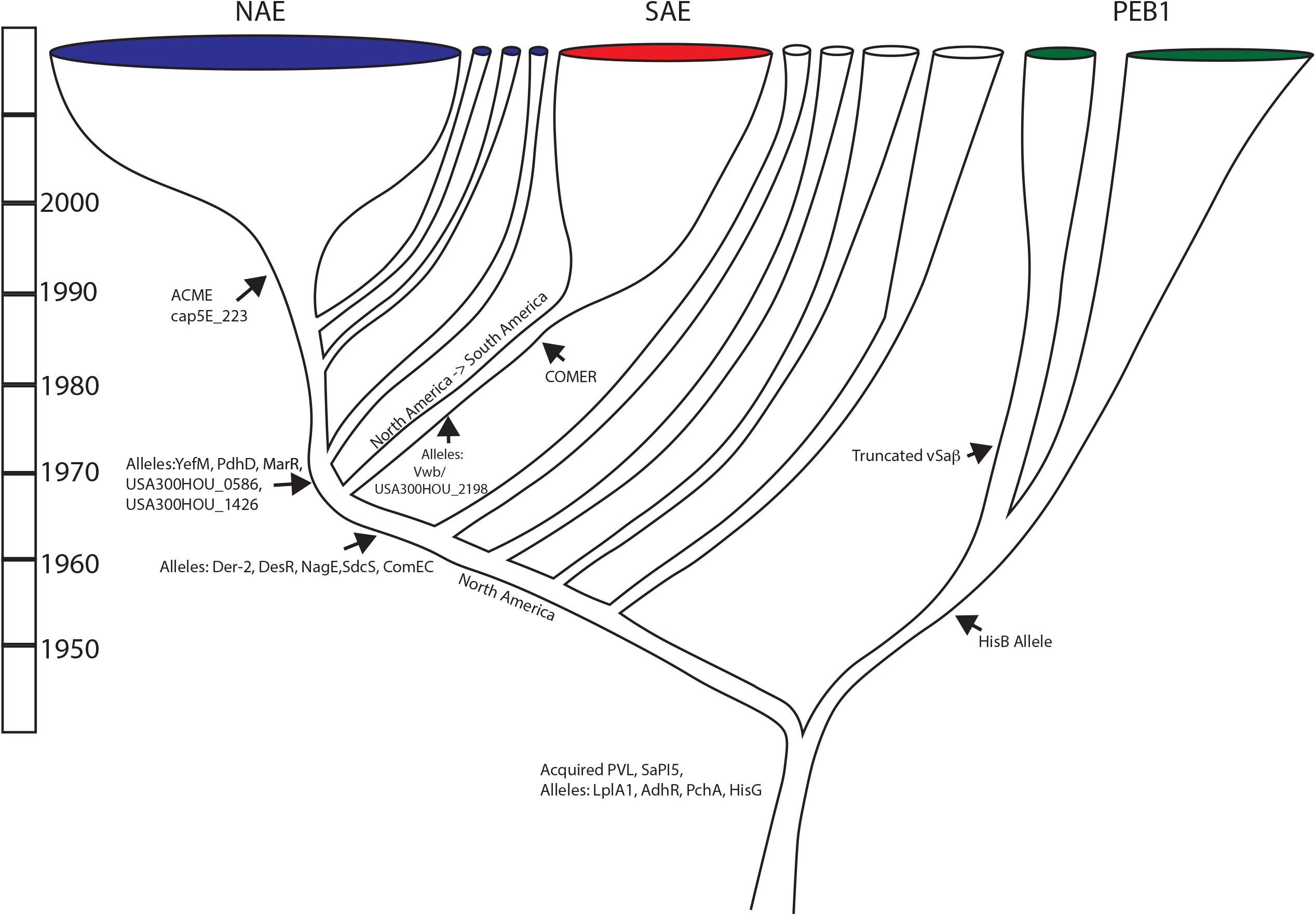
Evolutionary scenario of USA300: Summary of major evolution events of USA300 based on work presented. Major gene acquisition and allele changes are noted.

The first major divergence for this lineage separated the PEB1 clade from the clade that would go on to give rise to the NAE and SAE clades. The PEB1 clade was distinguished by a novel protein allele for HisB that may have affected its ability to make histidine. Within the PEB1 clade, an early divergence in the 1960s separated a group that would lose a large portion of the *v*Saβ locus. The PEB1 lineage spread globally and was transferred at least once, potentially twice, from North to South America. The sister lineage to PEB1 also underwent several transfer events from North America to South America during this time and after.

In the 1970s, the common ancestor of the epidemic lineages NAE and SAE arose, and it was characterized by novel protein alleles of Der_2, DesR, NagE, SdsC, and ComEC. With another North to South American introduction in the late 1960s or early 1970s this clade diverged into the epidemic lineages in North America and South America. It would be approximately 20 years later that the North American lineage would acquire the ACME element and the *cap5E*_223 mutation. The COMER locus was acquired in a separate event during the 1970s. The SAE lineage started to diversify in the 1980s, whereas most of the diversification that we see in the NAE lineage started in the 1990s. Thus, for both the NAE and the SAE clades, the acquisition of ACME and the COMER loci just precede epidemic spread, implicating these loci in transmissibility and/or fitness. The other genetic attributes that we have defined here, that trace this history may simply be neutral markers of evolutionary history, but they also may be changes that had functional impacts on the fitness of these lineages. As such, these genes and SNPs represent targets for future microbiological work that may lead to insights into the steps that lead to epidemic spread.

## Abbreviations

NAE: North American Epidemic clade
SAE: South American Epidemic clade
PEB: Pre-Epidemic clad
EB: Early Branching isolates
ACME: Arginine catabolic mobile element
COMER: Copper and mercury resistance mobile element
MSSA: Methicillin susceptible *Staphylococcus aureus*
MRSA: Methicillin resistant *Staphylococcus aureus*

## Acknowledgments

The authors thank Chanelle Ryan and Andries Feder for assistance with sequencing and the Planet and St. Geme labs for helpful discussions.

## References

2001. Methicillin-resistant Staphylococcus aureus skin or soft tissue infections in a state prison--Mississippi, 2000. MMWR Morb Mortal Wkly Rep, 50, 919–22.

Acker, K. P., Wong Fok Lung, T., West, E., Craft, J., Narechania, A., Smith, H., O’Brien, K., Moustafa, A. M., Lauren, C., Planet, P. J. & Prince, A. 2019. Strains of Staphylococcus aureus that Colonize and Infect Skin Harbor Mutations in Metabolic Genes. iScience, 19, 281–290.

Alvarez, C. A., Barrientes, O. J., Leal, A. L., Contreras, G. A., Barrero, L., Rincón, S., Diaz, L., Vanegas, N. & Arias, C. A. 2006. Community-associated methicillin-resistant Staphylococcus aureus, Colombia. Emerging infectious diseases, 12, 2000–2001.

Arias, C. A., Rincon, S., Chowdhury, S., Martínez, E., Coronell, W., Reyes, J., Nallapareddy, S. R. & Murray, B. E. 2008. MRSA USA300 clone and VREF—a U.S.-Colombian connection? N Engl J Med, 359, 2177–9.

Bosi, E., Monk, J. M., Aziz, R. K., Fondi, M., Nizet, V. & Palsson, B. Ø. 2016. Comparative genome-scale modelling of <i>Staphylococcus aureus</i> strains identifies strain-specific metabolic capabilities linked to pathogenicity. Proceedings of the National Academy of Sciences, 113, E3801–E3809.

Bouckaert, R., Vaughan, T. G., Barido-Sottani, J., Duchêne, S., Fourment, M., Gavryushkina, A., Heled, J., Jones, G., Kühnert, D., De Maio, N., Matschiner, M., Mendes, F. K., Müller, N. F., Ogilvie, H. A., Du Plessis, L., Popinga, A., Rambaut, A., Rasmussen, D., Siveroni, I., Suchard, M. A., Wu, C.-H., Xie, D., Zhang, C., Stadler, T. & Drummond, A. J. 2019. BEAST 2.5: An advanced software platform for Bayesian evolutionary analysis. PLOS Computational Biology, 15, e1006650.

Bowers, J. R., Driebe, E. M., Albrecht, V., Mcdougal, L. K., Granade, M., Roe, C. C., Lemmer, D., Rasheed, J. K., Engelthaler, D. M., Keim, P. & Limbago, B. M. 2018. Improved Subtyping of Staphylococcus aureus Clonal Complex 8 Strains Based on Whole-Genome Phylogenetic Analysis. mSphere, 3.

Boyle-Vavra, S., Li, X., Alam, M. T., Read, T. D., Sieth, J., Cywes-Bentley, C., Dobbins, G., David, M. Z., Kumar, N., Eells, S. J., Miller, L. G., Boxrud, D. J., Chambers, H. F., Lynfield, R., Lee, J. C., Daum, R. S. & Projan, S. J. 2015. USA300 and USA500 Clonal Lineages of Staphylococcus aureus Do Not Produce a Capsular Polysaccharide Due to Conserved Mutations in the cap5 Locus. mBio, 6, e02585–14.

Brynildsrud, O., Bohlin, J., Scheffer, L. & Eldholm, V. 2016. Rapid scoring of genes in microbial pan-genome-wide association studies with Scoary. Genome Biology, 17, 238.

Copin, R., Sause, W. E., Fulmer, Y., Balasubramanian, D., Dyzenhaus, S., Ahmed, J. M., Kumar, K., Lees, J., Stachel, A., Fisher, J. C., Drlica, K., Phillips, M., Weiser, J. N., Planet, P. J., Uhlemann, A. C., Altman, D. R., Sebra, R., Van Bakel, H., Lighter, J., Torres, V. J. & Shopsin, B. 2019. Sequential evolution of virulence and resistance during clonal spread of community-acquired methicillin-resistant Staphylococcus aureus. Proc Natl Acad Sci U S A, 116, 1745–1754.

Cordero, M., García-Fernández, J., Acosta, I. C., Yepes, A., Avendano-Ortiz, J., Lisowski, C., Oesterreicht, B., Ohlsen, K., Lopez-Collazo, E., Förstner, K. U., Eulalio, A. & Lopez, D. 2022. The induction of natural competence adapts staphylococcal metabolism to infection. Nature Communications, 13, 1525.

David, M. Z., Acree, M. E., Sieth, J. J., Boxrud, D. J., Dobbins, G., Lynfield, R., Boyle-Vavra, S., Daum, R. S. & Burnham, C.-A. D. 2015. Pediatric Staphylococcus aureus Isolate Genotypes and Infections from the Dawn of the Community-Associated Methicillin-Resistant S. aureus Epidemic Era in Chicago, 1994 to 1997. Journal of Clinical Microbiology, 53, 2486–2491.

Davis, J. J., Wattam, A. R., Aziz, R. K., Brettin, T., Butler, R., Butler, R. M., Chlenski, P., Conrad, N., Dickerman, A., Dietrich, E. M., Gabbard, J. L., Gerdes, S., Guard, A., Kenyon, R. W., Machi, D., Mao, C., Murphy-Olson, D., Nguyen, M., Nordberg, E. K., Olsen, G. J., Olson, R. D., Overbeek, J. C., Overbeek, R., Parrello, B., Pusch, G. D., Shukla, M., Thomas, C., Vanoeffelen, M., Vonstein, V., Warren, A. S., Xia, F., Xie, D., Yoo, H. & Stevens, R. 2019. The PATRIC Bioinformatics Resource Center: expanding data and analysis capabilities. Nucleic Acids Research, 48, D606–D612.

Dehouck, Y., Kwasigroch, J. M., Gilis, D. & Rooman, M. 2011. PoPMuSiC 2.1: a web server for the estimation of protein stability changes upon mutation and sequence optimality. BMC Bioinformatics, 12, 151.

Didelot, X. & Wilson, D. J. 2015. ClonalFrameML: Efficient Inference of Recombination in Whole Bacterial Genomes. PLOS Computational Biology, 11, e1004041.

Diep, B. A., Gill, S. R., Chang, R. F., Phan, T. H., Chen, J. H., Davidson, M. G., Lin, F., Lin, J., Carleton, H. A., Mongodin, E. F., Sensabaugh, G. F. & Perdreau-Remington, F. 2006. Complete genome sequence of USA300, an epidemic clone of community-acquired meticillin-resistant Staphylococcus aureus. The Lancet, 367, 731–739.

Dietl, A. M., Amich, J., Leal, S., Beckmann, N., Binder, U., Beilhack, A., Pearlman, E. & Haas, H. 2016. Histidine biosynthesis plays a crucial role in metal homeostasis and virulence of Aspergillus fumigatus. Virulence, 7, 465–76.

Fernández, P., Díaz, A. R., Ré, M. F., Porrini, L., De Mendoza, D., Albanesi, D. & Mansilla, M. C. 2020. Identification of Novel Thermosensors in Gram-Positive Pathogens. Frontiers in Molecular Biosciences, 7.

Fernández, P., Porrini, L., Albanesi, D., Abriata, L. A., Peraro, M. D., Mendoza, D. D., Mansilla, M. C. & Cossart, P. F. 2019. Transmembrane Prolines Mediate Signal Sensing and Decoding in Bacillus subtilis DesK Histidine Kinase. mBio, 10, e02564–19.

Gabryszewski, S. J., Wong Fok Lung, T., Annavajhala, M. K., Tomlinson, K. L., Riquelme, S. A., Khan, I. N., Noguera, L. P., Wickersham, M., Zhao, A., Mulenos, A. M., Peaper, D., Koff, J. L., Uhlemann, A. C. & Prince, A. 2019. Metabolic Adaptation in Methicillin-Resistant Staphylococcus aureus Pneumonia. Am J Respir Cell Mol Biol, 61, 185–197.

Hall, J. A. & Pajor, A. M. 2007. Functional reconstitution of SdcS, a Na+-coupled dicarboxylate carrier protein from Staphylococcus aureus. J Bacteriol, 189, 880–5.

Hasegawa, M., Kishino, H. & Yano, T.-A. 1985. Dating of the human-ape splitting by a molecular clock of mitochondrial DNA. Journal of Molecular Evolution, 22, 160–174.

Hwang, J., Tseitin, V., Ramnarayan, K., Shenderovich, M. D. & Inouye, M. 2012. Structure-based design and screening of inhibitors for an essential bacterial GTPase, Der. The Journal of Antibiotics, 65, 237–243.

Ishikawa, S. A., Zhukova, A., Iwasaki, W. & Gascuel, O. 2019. A Fast Likelihood Method to Reconstruct and Visualize Ancestral Scenarios. Molecular Biology and Evolution, 36, 2069–2085.

Johnson, M., Zaretskaya, I., Raytselis, Y., Merezhuk, Y., Mcginnis, S. & Madden, T. L. 2008. NCBI BLAST: a better web interface. Nucleic Acids Res, 36, W5–9.

Kaya, H., Hasman, H., Larsen, J., Stegger, M., Johannesen, T. B., Allesøe, R. L., Lemvigh, C. K., Aarestrup, F. M., Lund, O., Larsen, A. R. & Limbago, B. M. 2018. SCCmec Finder, a Web-Based Tool for Typing of Staphylococcal Cassette Chromosome mec in Staphylococcus aureus Using Whole-Genome Sequence Data. mSphere, 3, e00612–17.

Kennedy, A. D., Otto, M., Braughton, K. R., Whitney, A. R., Chen, L., Mathema, B., Mediavilla, J. R., Byrne, K. A., Parkins, L. D., Tenover, F. C., Kreiswirth, B. N., Musser, J. M. & Deleo, F. R. 2008. Epidemic community-associated methicillin-resistant Staphylococcus aureus: recent clonal expansion and diversification. Proc Natl Acad Sci U S A, 105, 1327–32.

King, M. D., Humphrey, B. J., Wang, Y. F., Kourbatova, E. V., Ray, S. M. & Blumberg, H. M. 2006. Emergence of community-acquired methicillin-resistant Staphylococcus aureus USA 300 clone as the predominant cause of skin and soft-tissue infections. Ann Intern Med, 144, 309–17.

Kläui, A. J., Boss, R. & Graber, H. U. 2019. Characterization and Comparative Analysis of the Staphylococcus aureus Genomic Island vSaβ: an In Silico Approach. Journal of bacteriology, 201, e00777–18.

Kwong, J. S. Torsten 2019. maskrc-svg,Masks recombinant regions in an alignment based on ClonalFrameML or Gubbins output.

Lepuschitz, S., Huhulescu, S., Hyden, P., Springer, B., Rattei, T., Allerberger, F., Mach, R. L. & Ruppitsch, W. 2018. Characterization of a community-acquired-MRSA USA300 isolate from a river sample in Austria and whole genome sequence based comparison to a diverse collection of USA300 isolates. Scientific reports, 8, 9467–9467.

Letunic, I. & Bork, P. 2021. Interactive Tree Of Life (iTOL) v5: an online tool for phylogenetic tree display and annotation. Nucleic Acids Research, 49, W293–W296.

Maddison, W. P. & Maddison, D. R. 2021. Mesquite: a modular system for evolutionary analysis.

Martin Simonsen, T. M., CHRISTIAN N. S. Pedersen 2008. Rapid Neighbour Joining. Proceedings of the 8th Workshop in Algorithms in Bioinformatics (WABI), LNBI 5251, 113–122.

Mohamed, N., Timofeyeva, Y., Jamrozy, D., Rojas, E., Hao, L., Silmon De Monerri, N. C., Hawkins, J., Singh, G., Cai, B., Liberator, P., Sebastian, S., Donald, R. G. K., Scully, I. L., Jones, C. H., Creech, C. B., Thomsen, I., Parkhill, J., Peacock, S. J., Jansen, K. U., Holden, M. T. G. & Anderson, A. S. 2019. Molecular epidemiology and expression of capsular polysaccharides in Staphylococcus aureus clinical isolates in the United States. PLOS ONE, 14, e0208356.

Morikawa, K., Takemura, A. J., Inose, Y., Tsai, M., Nguyen Thi, L. T., Ohta, T. & Msadek, T. 2012. Expression of a Cryptic Secondary Sigma Factor Gene Unveils Natural Competence for DNA Transformation in Staphylococcus aureus. PLOS Pathogens, 8, e1003003.

Moustafa, A. M. & Planet, P. J. 2020. WhatsGNU: a tool for identifying proteomic novelty. Genome Biology, 21, 58.

Ng, P. C. & Henikoff, S. 2003. SIFT: Predicting amino acid changes that affect protein function. Nucleic acids research, 31, 3812–3814.

Ondov, B. D., Starrett, G. J., Sappington, A., Kostic, A., Koren, S., Buck, C. B. & Phillippy, A. M. 2019. Mash Screen: high-throughput sequence containment estimation for genome discovery. Genome Biology, 20, 232.

Page, A. J., Cummins, C. A., Hunt, M., Wong, V. K., Reuter, S., Holden, M. T. G., Fookes, M., Falush, D., Keane, J. A. & Parkhill, J. 2015. Roary: rapid large-scale prokaryote pan genome analysis. Bioinformatics, 31, 3691–3693.

Page, A. J., Taylor, B., Delaney, A. J., Soares, J., Seemann, T., Keane, J. A. & Harris, S. R. 2016. SNP-sites: rapid efficient extraction of SNPs from multi-FASTA alignments. Microb Genom, 2, e000056.

Patrick, W. M., Quandt, E. M., Swartzlander, D. B. & Matsumura, I. 2007. Multicopy suppression underpins metabolic evolvability. Mol Biol Evol, 24, 2716–22.

Petit, R. A., 3rd & Read, T. D. 2018. Staphylococcus aureus viewed from the perspective of 40,000+ genomes. PeerJ, 6, e5261–e5261.

Pimentel, Z. T. & Zhang, Y. 2018. Evolution of the Natural Transformation Protein, ComEC, in Bacteria. Frontiers in Microbiology, 9.

Planet, P. J., Diaz, L., Kolokotronis, S. O., Narechania, A., Reyes, J., Xing, G., Rincon, S., Smith, H., Panesso, D., Ryan, C., Smith, D. P., Guzman, M., Zurita, J., Sebra, R., Deikus, G., Nolan, R. L., Tenover, F. C., Weinstock, G. M., Robinson, D. A. & Arias, C. A. 2015. Parallel Epidemics of Community-Associated Methicillin-Resistant Staphylococcus aureus USA300 Infection in North and South America. J Infect Dis, 212, 1874–82.

Planet, P. J., Diaz, L., Rios, R. & Arias, C. A. 2016. Global Spread of the Community-Associated Methicillin-Resistant Staphylococcus aureus USA300 Latin American Variant. J Infect Dis, 214, 1609–1610.

Planet, P. J., Larussa, S. J., Dana, A., Smith, H., Xu, A., Ryan, C., Uhlemann, A. C., Boundy, S., Goldberg, J., Narechania, A., Kulkarni, R., Ratner, A. J., Geoghegan, J. A., Kolokotronis, S. O. & Prince, A. 2013. Emergence of the epidemic methicillin-resistant Staphylococcus aureus strain USA300 coincides with horizontal transfer of the arginine catabolic mobile element and speG-mediated adaptations for survival on skin. mBio, 4, e00889–13.

Reyes, J., Rincón, S., Díaz, L., Panesso, D., Contreras, G. A., Zurita, J., Carrillo, C., Rizzi, A., Guzmán, M., Adachi, J., Chowdhury, S., Murray, B. E. & Arias, C. A. 2009. Dissemination of methicillin-resistant Staphylococcus aureus USA300 sequence type 8 lineage in Latin America. Clin Infect Dis, 49, 1861–7.

Roberts, R. B., De Lencastre, A., Eisner, W., Severina, E. P., Shopsin, B., Kreiswirth, B. N. & Tomasz, A. 1998. Molecular epidemiology of methicillin-resistant Staphylococcus aureus in 12 New York hospitals. MRSA Collaborative Study Group. J Infect Dis, 178, 164–71.

Seemann, T. 2014. Prokka: rapid prokaryotic genome annotation. Bioinformatics, 30, 2068–2069.

Seybold, U., Kourbatova, E. V., Johnson, J. G., Halvosa, S. J., Wang, Y. F., King, M. D., Ray, S. M. & Blumberg, H. M. 2006. Emergence of Community-Associated Methicillin-Resistant Staphylococcus aureus USA300 Genotype as a Major Cause of Health Care—Associated Blood Stream Infections. Clinical Infectious Diseases, 42, 647–656.

Stamatakis, A. 2014. RAxML version 8: a tool for phylogenetic analysis and post-analysis of large phylogenies. Bioinformatics, 30, 1312–1313.

Straus, L., Stegger, M., Akpaka, P. E., Alabi, A., Breurec, S., Coombs, G., Egyir, B., Larsen, A. R., Laurent, F., Monecke, S., Peters, G., Skov, R., Strommenger, B., Vandenesch, F., Schaumburg, F. & Mellmann, A. 2017. Origin, evolution, and global transmission of community-acquired Staphylococcus aureus ST8. Proceedings of the National Academy of Sciences, 114, E10596–E10604.

Swofford, D. L. 2003. PAUP*. Phylogenetic Analysis Using Parsimony (*and Other Methods). Sinauer Associates, Sunderland, Massachusetts., Version 4.

Uhlemann, A. C., Dordel, J., Knox, J. R., Raven, K. E., Parkhill, J., Holden, M. T., Peacock, S. J. & Lowy, F. D. 2014. Molecular tracing of the emergence, diversification, and transmission of S. aureus sequence type 8 in a New York community. Proc Natl Acad Sci U S A, 111, 6738–43.

Villanueva, R. A. M. & Chen, Z. J. 2019. ggplot2: Elegant Graphics for Data Analysis (2nd ed.). Measurement: Interdisciplinary Research and Perspectives, 17, 160–167.

Von Dach, E., Diene, S. M., Fankhauser, C., Schrenzel, J., Harbarth, S. & François, P. 2016. Comparative Genomics of Community-Associated Methicillin-Resistant Staphylococcus aureus Shows the Emergence of Clone ST8-USA300 in Geneva, Switzerland. J Infect Dis, 213, 1370–9.

Wick, R. R., Judd, L. M., Gorrie, C. L. & Holt, K. E. 2017. Unicycler: Resolving bacterial genome assemblies from short and long sequencing reads. PLOS Computational Biology, 13, e1005595.

Xiao, X., Li, Y., Li, L. & Xiong, Y. Q. 2022. Identification of Methicillin-Resistant Staphylococcus aureus (MRSA) Genetic Factors Involved in Human Endothelial Cells Damage, an Important Phenotype Correlated with Persistent Endovascular Infection. Antibiotics, 11, 316.

Yu, G. 2020. Using ggtree to Visualize Data on Tree-Like Structures. Curr Protoc Bioinformatics, 69, e96.

Zhou, Y.-H., Xu, C.-G., Yang, Y.-B., Xing, X.-X., Liu, X., Qu, Q.-W., Ding, W.-Y., Bello-Onaghise, G. S. & Li, Y.-H. 2018. Histidine Metabolism and IGPD Play a Key Role in Cefquinome Inhibiting Biofilm Formation of Staphylococcus xylosus. Frontiers in Microbiology, 9.

