## Supplemental Figures for "Pre-epidemic evolution of the USA300 clade and a molecular key for classification"

**Supplemental: 5 tables and 8 figures**

**Table S1**: Genome quality statistics for newly reported genomes generated using QUAST.

| | **Assembly** | | | |
|---|---|---|---|---|
| **Statistic** | **2m-n** | **BK2448** | **BK2561B** | **65_669** |
| # contigs (>= 0 bp) | 75 | 64 | 77 | 26 |
| # contigs (>= 1000 bp) | 38 | 28 | 34 | 19 |
| # contigs (>= 5000 bp) | 32 | 24 | 26 | 12 |
| # contigs (>= 10000 bp) | 25 | 18 | 23 | 11 |
| # contigs (>= 25000 bp) | 19 | 15 | 20 | 10 |
| # contigs (>= 50000 bp) | 16 | 12 | 17 | 9 |
| Total length (>= 0 bp) | 2899456 | 2874324 | 2871754 | 2777101 |
| Total length (>= 1000 bp) | 2892994 | 2867581 | 2864850 | 2775911 |
| Total length (>= 5000 bp) | 2867348 | 2852203 | 2838817 | 2753096 |
| Total length (>= 10000 bp) | 2827327 | 2814495 | 2815656 | 2747552 |
| Total length (>= 25000 bp) | 2718877 | 2760373 | 2753311 | 2726838 |
| Total length (>= 50000 bp) | 2605280 | 2643903 | 2646126 | 2684464 |
| # contigs | 38 | 30 | 34 | 19 |
| Largest contig | 343261 | 610876 | 677524 | 883770 |
| Total length | 2892994 | 2869164 | 2864850 | 2775911 |
| GC (%) | 32.86 | 32.87 | 32.73 | 32.86 |
| N50 | 198538 | 300945 | 202206 | 566023 |
| N75 | 112194 | 147425 | 113145 | 348456 |
| L50 | 5 | 4 | 4 | 2 |
| L75 | 11 | 7 | 9 | 4 |
| # N's per 100 kbp | 20.81 | 19.87 | 7.54 | 4.29 |
| All statistics are based on contigs of size >= 500 bp, unless otherwise noted (e.g., "# contigs (>= 0 bp)" and "Total length (>= 0 bp)" include all contigs). | | | | |

**Table S2**: Characterization of *cap5D* and *cap5D* mutations

| Allele | Mutation (nt) | Mutation Nature (AA/ protein) | Active Enzyme? |
|---|---|---|---|
| cap5E_223 | G223T | Asp75Tyr in enzyme active site | No |
| cap5E_223_A170G | A170G, G223T | Tyr57Cys, Asp75Tyr in enzyme active site | No |
| cap5E_223_T17C | T17C, G223T | Lle6Thr, Asp75Tyr in enzyme active site | No |
| cap5E_223_G117A | G117A, G223T | silent, Asp75Tyr in enzyme active site | No |
| cap5E_WT | None | wt | Yes |
| cap5E_T116C | T166C | Leu58Phe | ? |
| cap5E_223_G268C | G223T, G268C | Asp75Tyrin enzyme active site, Glu88Gln | No |
| cap5E_A85G | A85G | Lle28Val | ? |
| cap5E_T458C | T458C | silent- wt | Yes |
| cap5E_G613A | G613A | Ala205Thr | ? |
| cap5E_A137**Δ** | A137delete | Frameshift | No (?) |
| cap5E_518AG | 518AG | Frameshift | No (?) |
| cap5D_994AA | 994AA | Frameshift, truncated Cap5D | No |
| cap5D_994AA_C737T | C737T, 994AA | Frameshift, truncated Cap5D | No |
| cap5D_994AA_G415A | G415A, 994AA | Frameshift, truncated Cap5D | No |
| cap5D_994AA_G555A | G555A, 994AA | Frameshift, truncated Cap5D | No |
| capDE_WT | None | wt | Yes |
| cap5D_**Δ**210 nt | Missing 210nt | no protein made | No (?) |
| no *cap5D* found | NA | NA | No (?) |

**Table S3**: Novel protein alleles specific to USA300-PEB1 clade. *mutation from MRCA with EB/SAE/NAE clades to PEB1

| Gene | Protein | Mutation* (nt) | Mutation* Nature (AA/ protein) | Mutation Tolerated_SIFT | Mutation Tolerated_dezyme | PBD (WT) |
|---|---|---|---|---|---|---|
| *hisB* | Imidazoleglycerol-phosphate dehydratase | C158G | Ala53Gly | No | No | 2AE8 |

**Table S4**: Novel protein alleles specific to USA300-SAE and NAE clades. *=mutation from MRCA with non-epidemic genomes.

| Gene | Protein | Mutation* (nt) | Mutation* Nature (AA/ protein) | Mutation Tolerated_SIFT |
|---|---|---|---|---|
| *der* | GTPase Der | A1094G | D365G | Yes |
| *desR* | Transcriptional regulatory protein DesR | A154T | I52F | Yes |
| *nagE* | PTS system N-acetylglucosamine-specific EIICBA component | C224T | A75V | Yes |
| *comEC* | Natural transformation protein | G608T | G203V | No |
| *sdcS* | Sodium-dependent dicarboxylate transporter SdcS | C712T | H238Y | No |

**Table S5: Basis for deriving sensitivity and specificity.** Table showing positive and negative values for A. USA300 or B. NAE. MK= molecular key

## A

| | USA300 on Tree | Not USA300 on Tree |
|---|---|---|
| USA300 on MK | True Postive: 4,069 | False Positive: 1 |
| Not USA300 on MK | False Negative: 7 | True Negative: 38,732 |

## B

| | NAE on Tree | Not NAE on Tree |
|---|---|---|
| NAE on MK | True Postive: 3,945 | False Positive: 24 |
| Not NAE on MK | False Negative: 26 | True Negative: 38,832 |

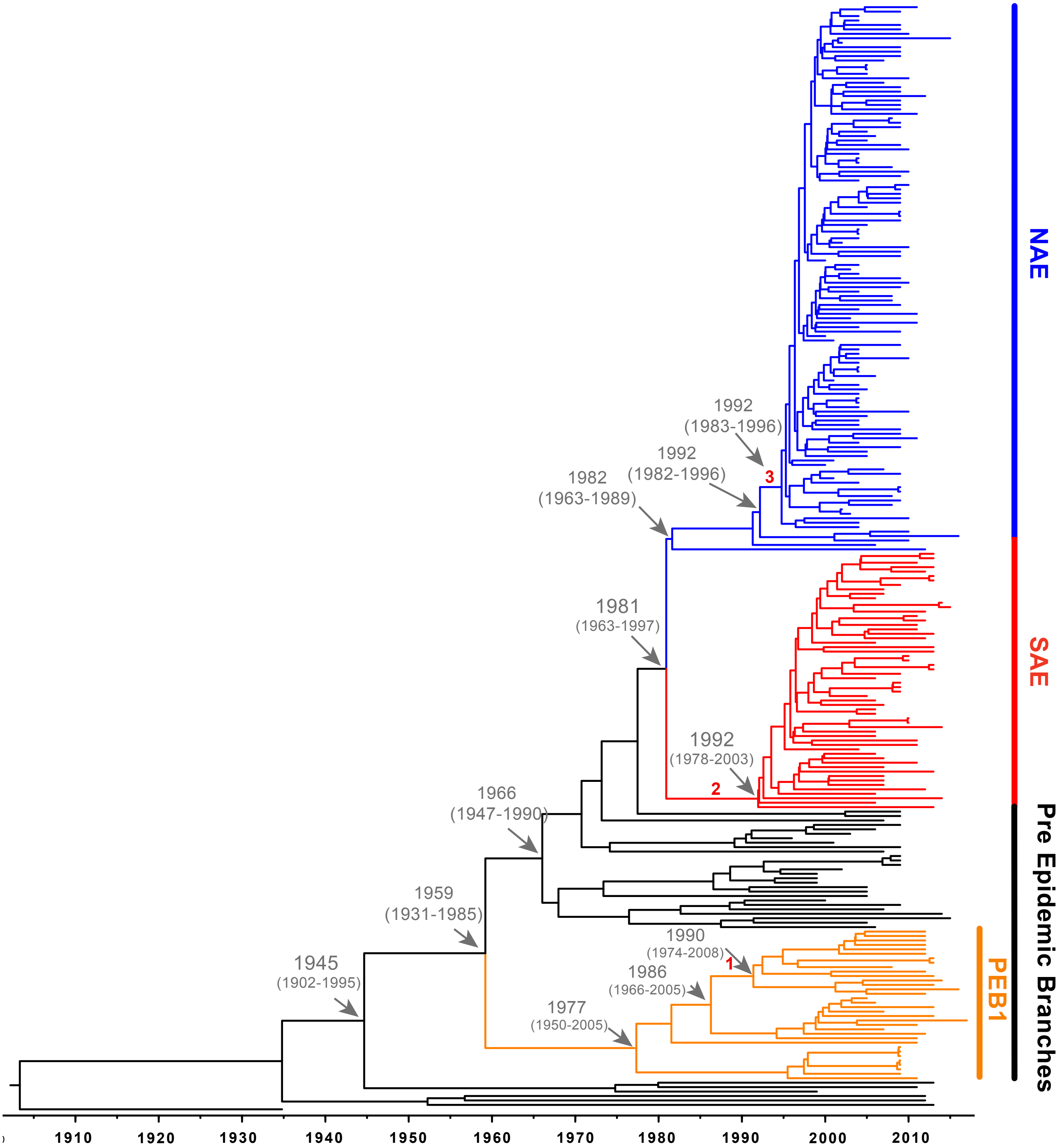


**Figure S1. Bayesian maximum clade tree calculated from 305332 sampled trees using a relaxed clock.** Tree generated using uncorrelated lognormal relaxed clock and a constant-size coalescent population. Four independent runs of 200 million MCMC steps with a 5,000-step thinning were completed. For each run, 10% of the first posterior samples were removed as a burn-in and the posterior estimates from 4 individual runs were combined to form a posterior sample density based on 760 million samples. Important MRCA are indicated as dates (95% HPD) and introduction of key genetic elements are labelled (1. vSaß loss, 2. COMER acquisition, 3. ACME acquisition)

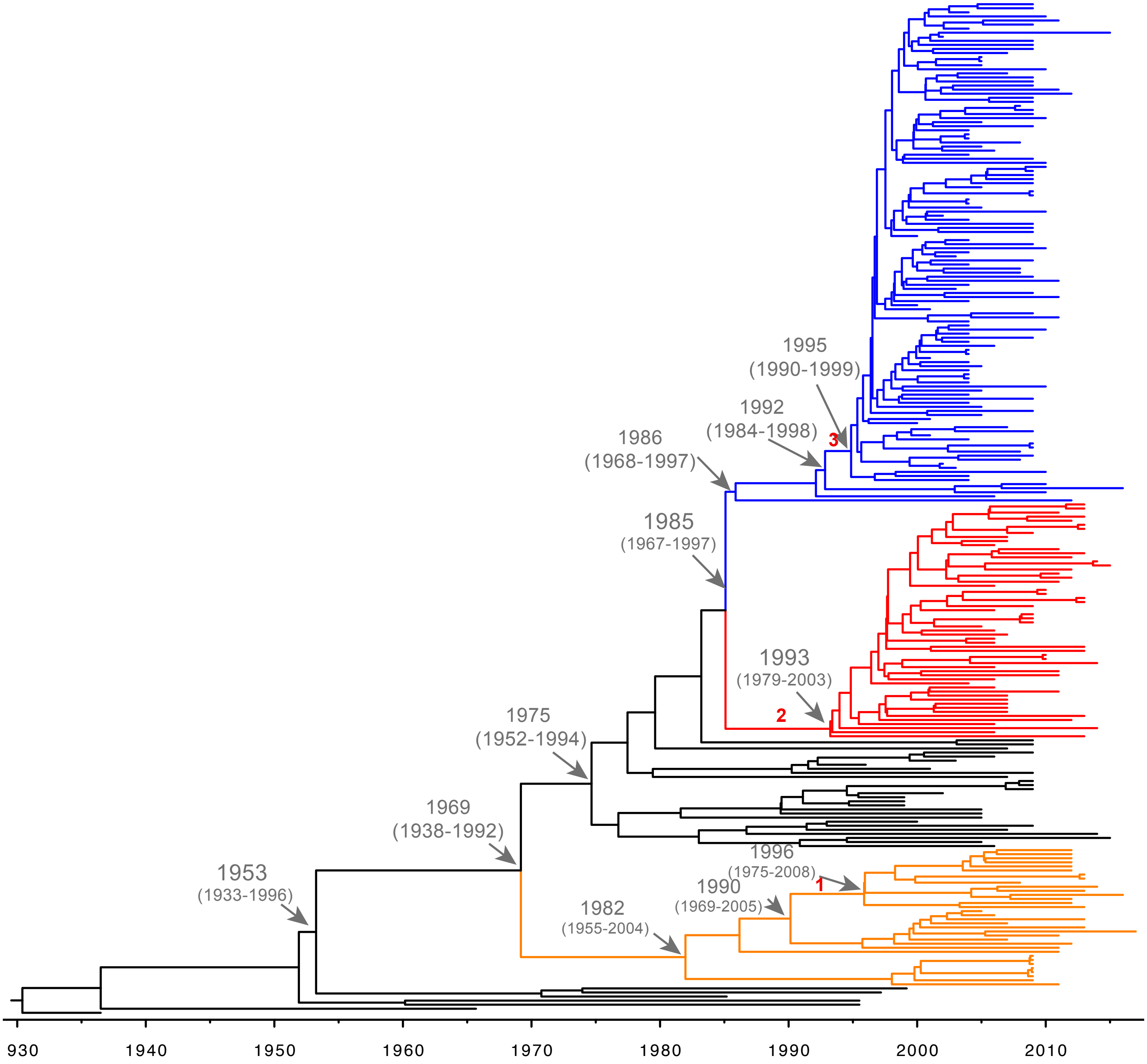


**Figure S2. Bayesian maximum clade tree calculated from 148936 sampled trees using a relaxed clock.** Tree generated using uncorrelated lognormal relaxed clock and a coalescent exponential population. Two independent runs of 200 million MCMC steps with a 5,000-step thinning were completed. For each run, 10% of the first posterior samples were removed as a burn-in and the posterior estimates from 2 individual runs were combined to form a posterior sample density based on 380 million samples. Important MRCA are indicated as dates (95% HPD) and introduction of key genetic elements are labelled (1. vSaß loss, 2. COMER acquisition, 3. ACME acquisition)

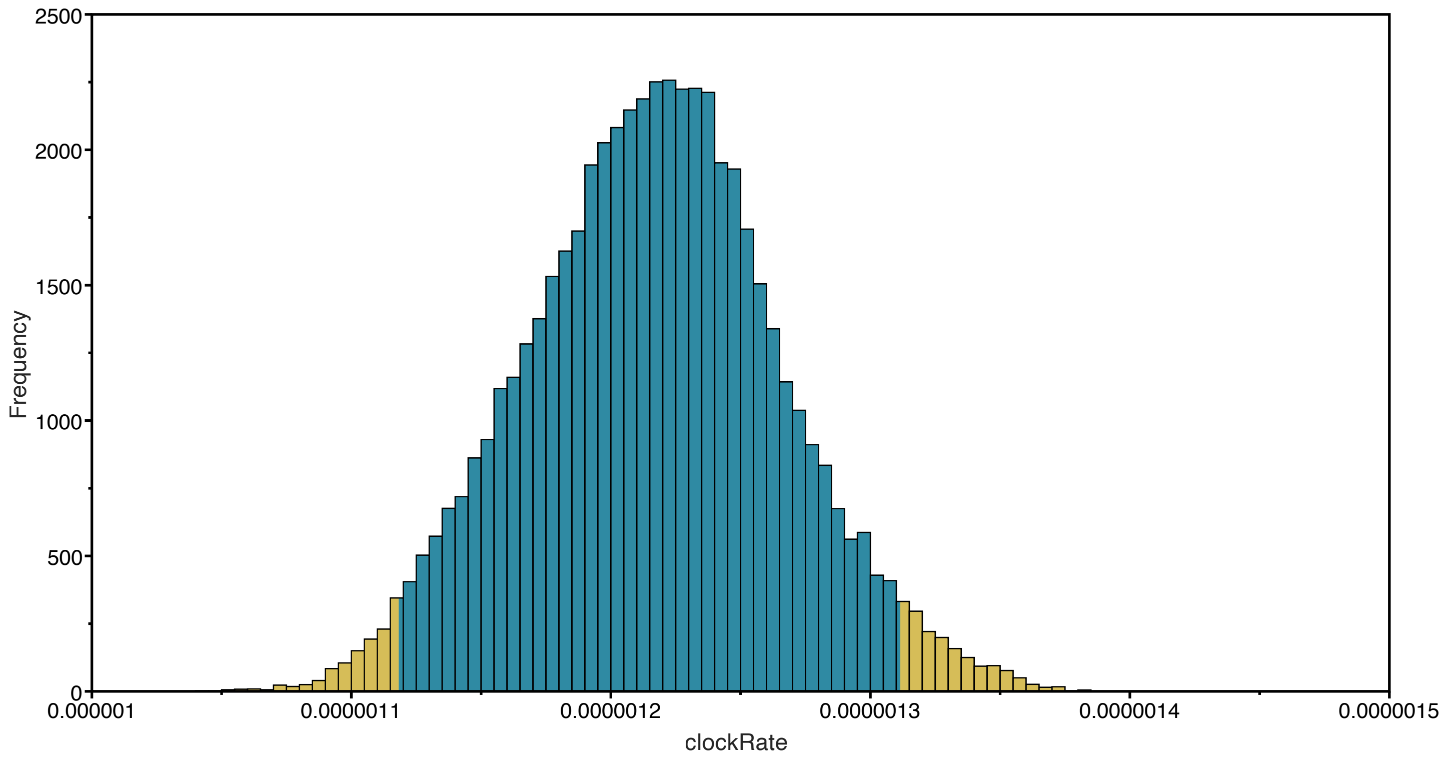


**Figure S3.** Mean evolutionary rate (substitutions/site/year) across the whole phylogenetic tree using a strict clock model with a random starting tree and a coalescent constant population. Mean=1.2164E-6, Median=1.217E-6, SE=4.6231E-9, SD=4.8944E-8, 95% HPD Interval=1.1184E-6, 1.3117E-6.

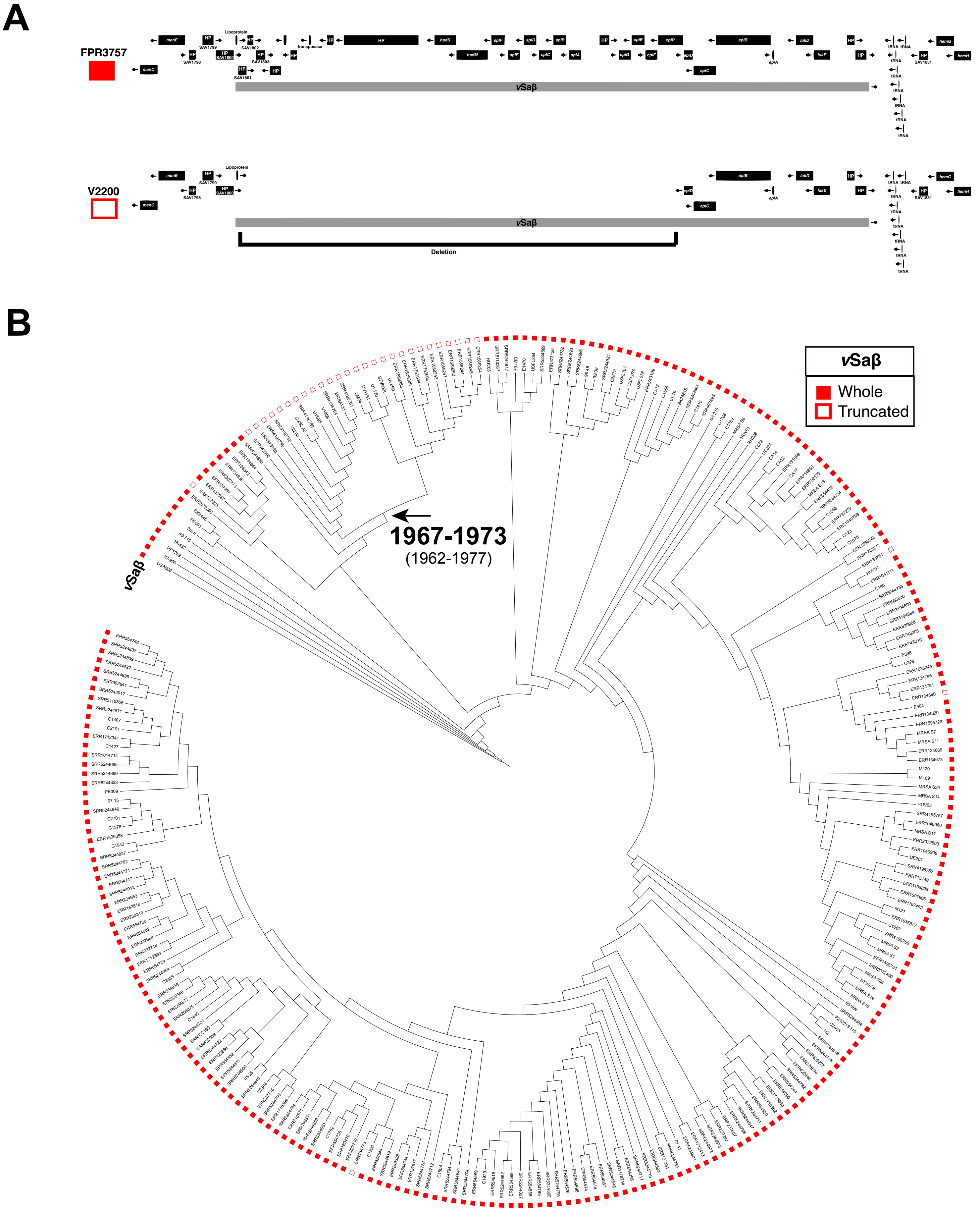


**Figure S4: Majority of genomes in the PEB1 are missing part of the genomic island vSaβ, most likely through an excision event**. (A) Representative alignment of whole (FPR3757) or truncated (V2200) genomic island vSaβ. Arrows show orientation of open reading frames. HP= hypothetical protein. (B) Phylogenetic analysis of 276 S. aureus strains showing the presence or absence of whole vSaβ in USA300. Estimated date of excision is indicated (95% HPD).

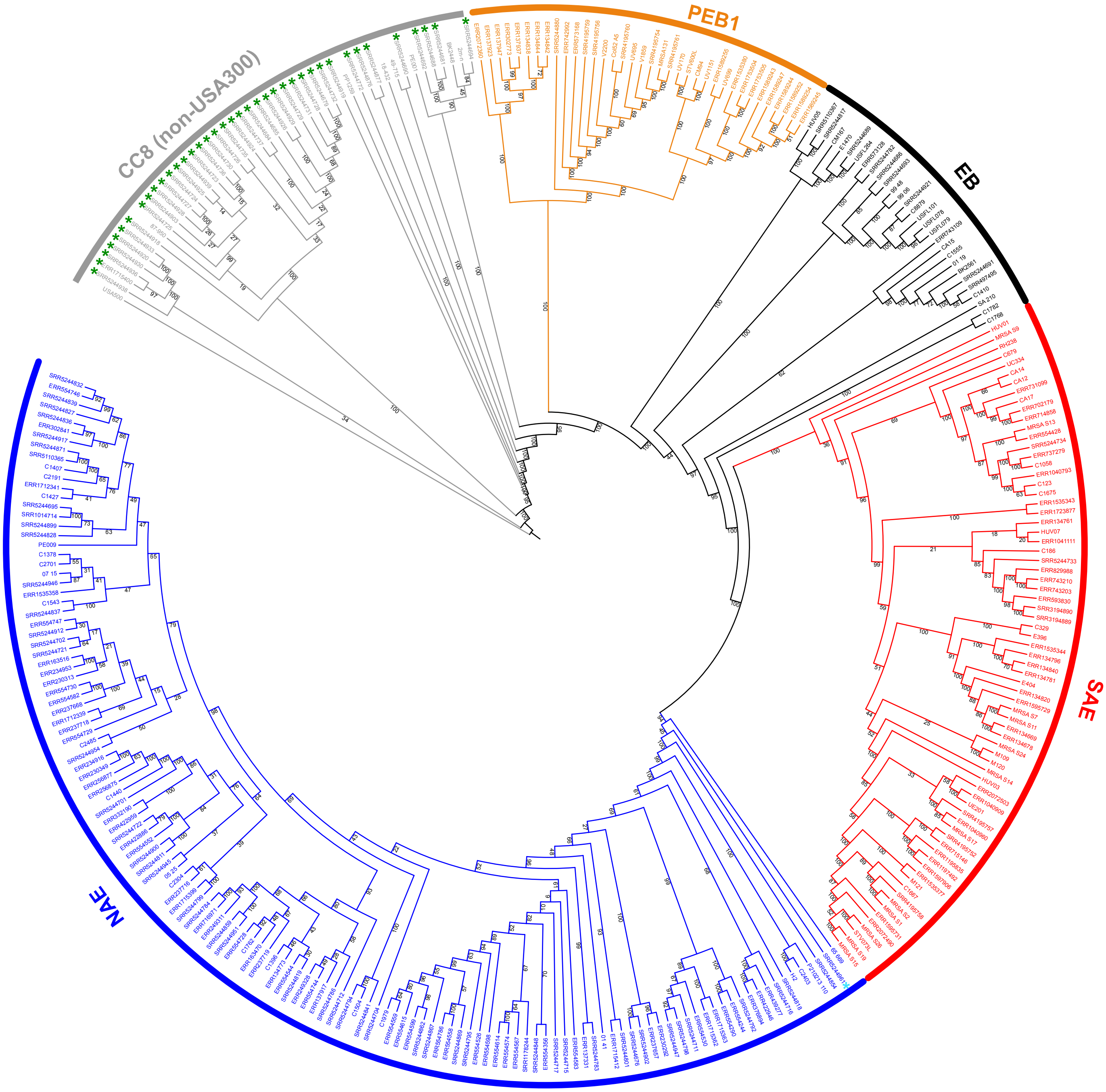


**Figure S5: Phylogenetic analysis of USA300 and non-USA300 CC8 genomes:** Branches and names are colored according to clade. Additional CC8 genomes in this analysis are marked with a green asterisk and the added "SRR5244961" genome is marked with cyan asterisk.

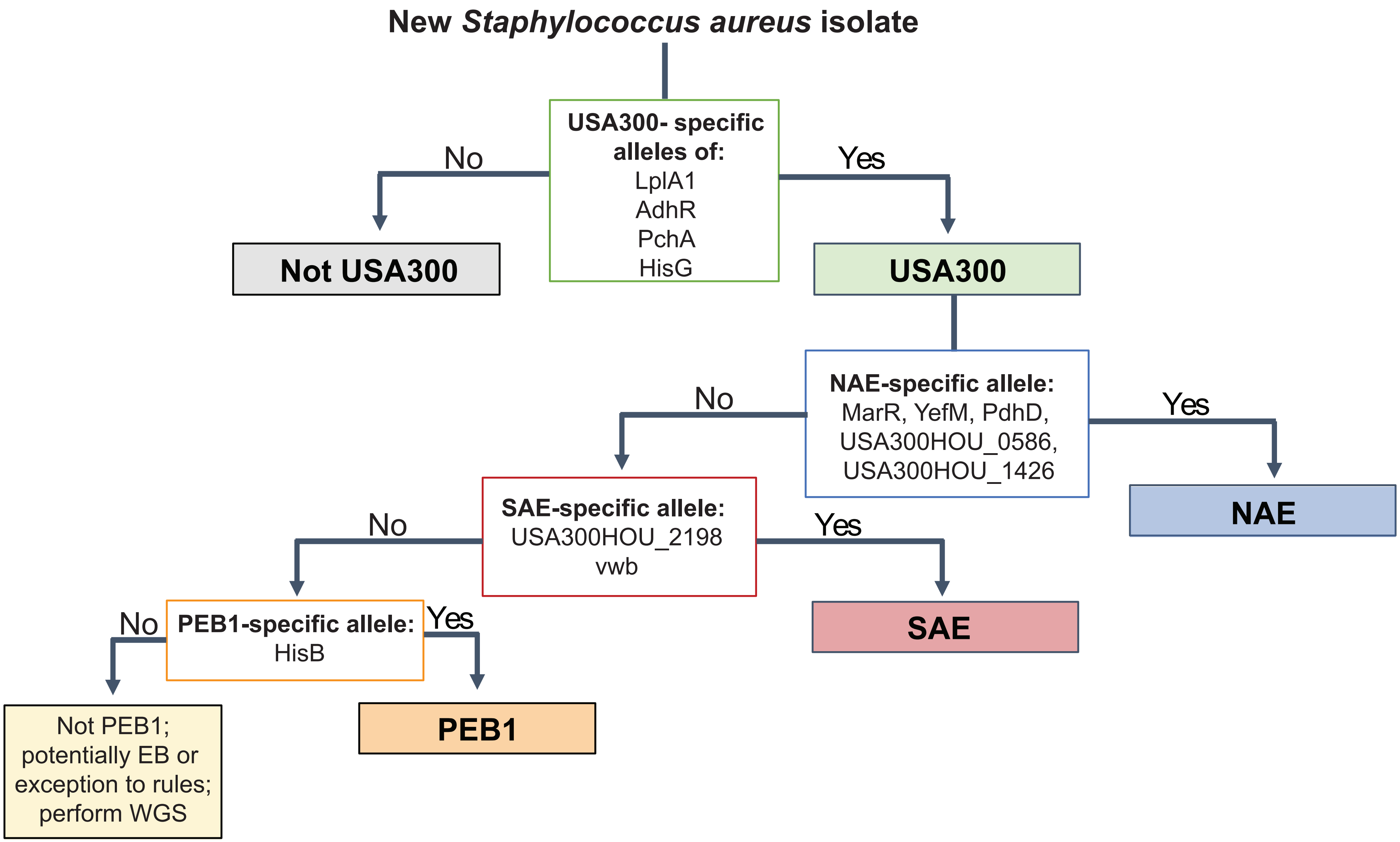


**Figure S6: Suggested process for identifying a newly isolated *Staphylococcus aureus*.** Alleles are specific to each clade and do not need to be used in conjunction with a prerequisite. For example, you can test for PEB1 without first determining that your isolate is not NAE. The order suggested reflects the probability that a new isolate will most likely be NAE, then SAE, and finally PEB1.

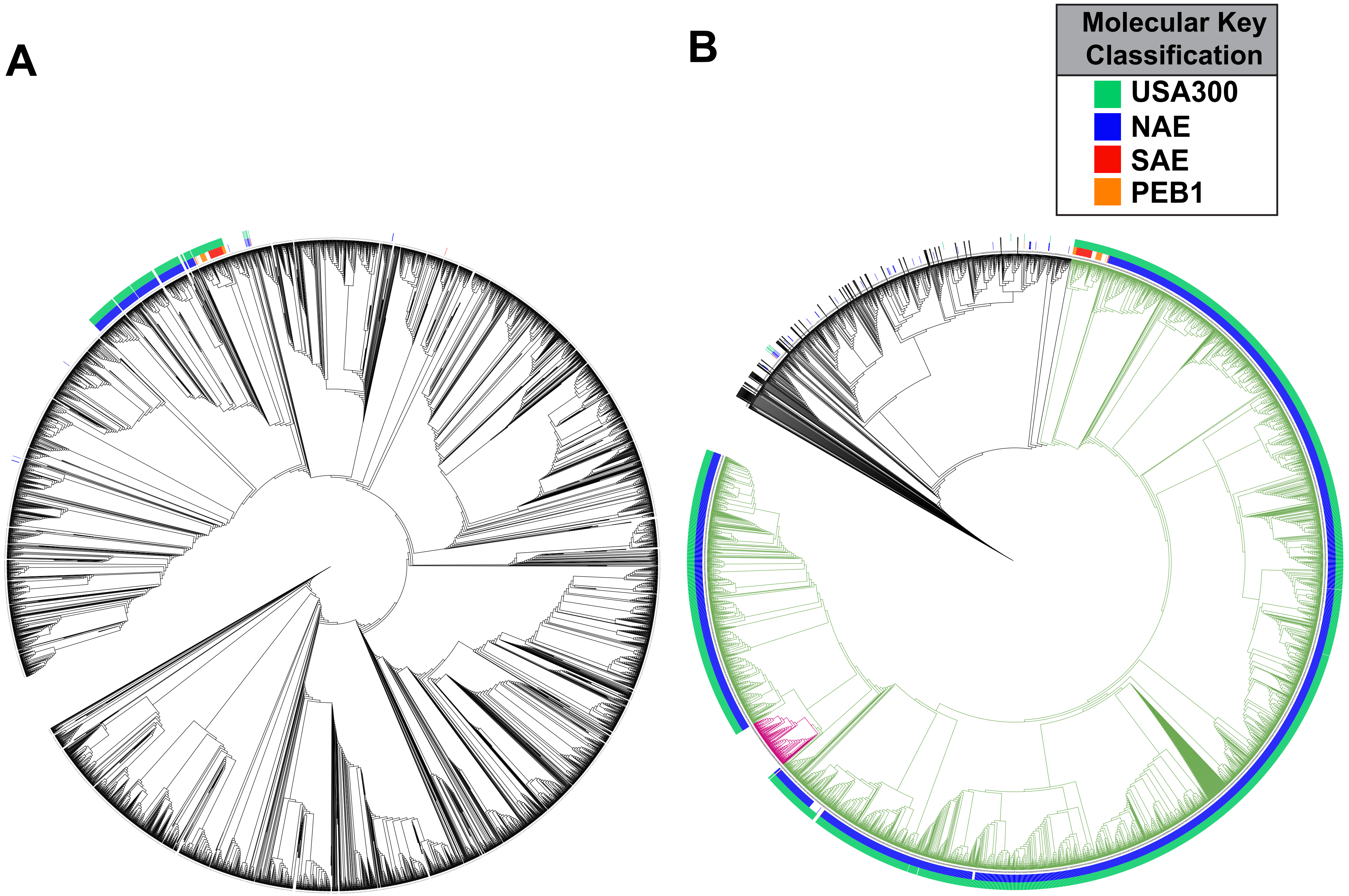


**Figure S7: Neighbor joining tree of all 42949 S. aureus genomes on Staphopia.** Genomes classified as USA300 using the allele-based molecular key are labeled with green (inner ring) and genomes possessing clade-specific alleles are colored according to alleles used (outer ring). A. Is the fully expanded tree. B. is the same tree with all clades besides the putative USA300 clade (green branches) collapsed. The clade in located within USA300 but not identified as USA300 using the molecular key is labeled in pink.

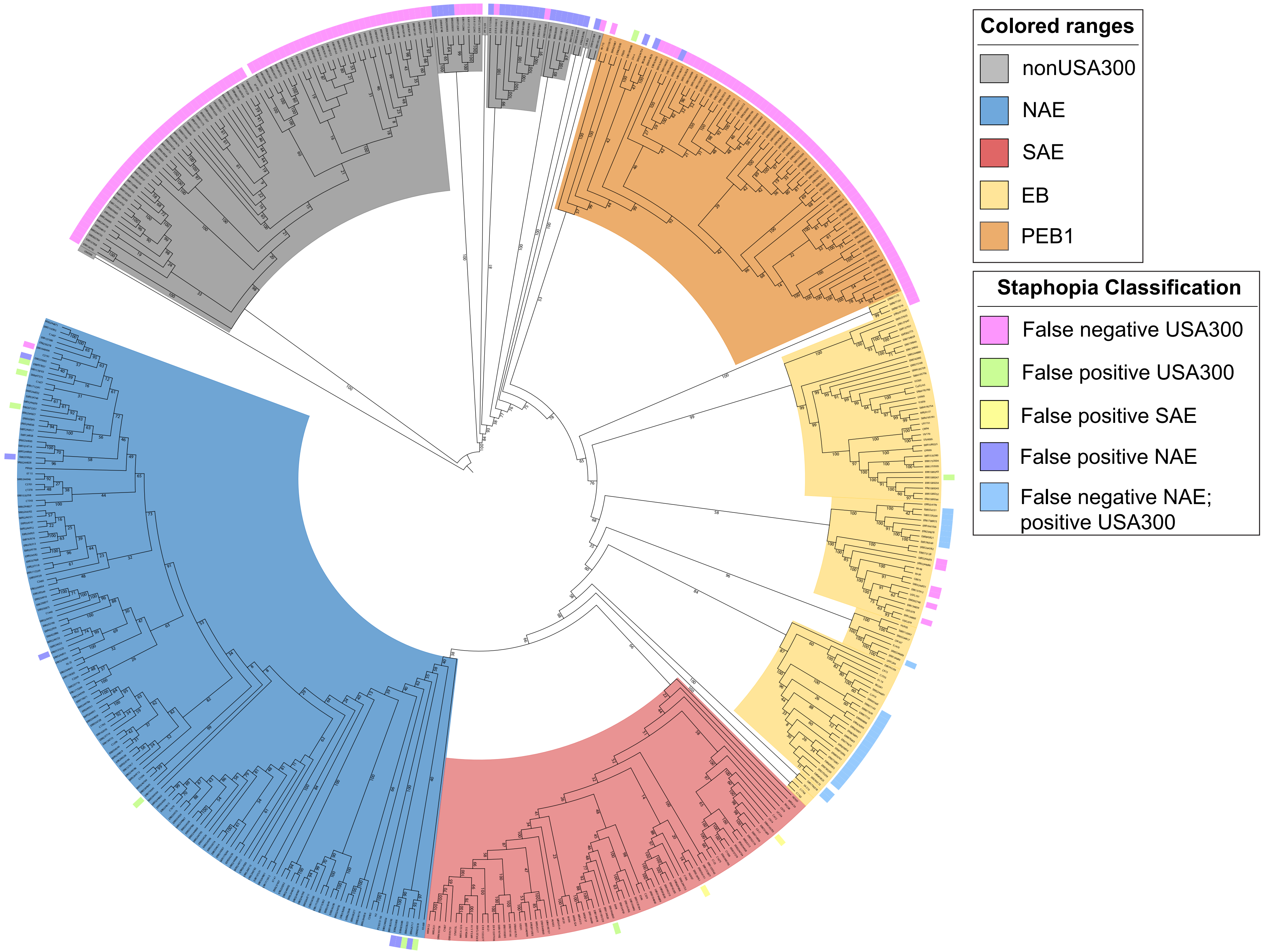


**Figure S8. Maximum likelihood tree with added conflicting *S. aureus* genomes:** Genomes for which the classification using the allelic molecular key were discordant with the relationships based on the NJ Staphopia tree were re-assessed using maximum likelihood in combination with 276 genomes from Fig. 1. The branches are colored blue for NAE, red for SAE, yellow for early branching clades and PEB1, and gray for non-USA300 CC8. The outer ring color indicates the results of the initial analysis based on classification using the allelic molecular key as the "experiment" and Staphopia tree as "true."
